# Supervised biological network alignment with graph neural networks

**DOI:** 10.1101/2023.04.24.538184

**Authors:** Kerr Ding, Sheng Wang, Yunan Luo

## Abstract

**Motivation:** Despite the advances in sequencing technology, massive proteins with known sequences remain functionally unannotated. Biological network alignment (NA), which aims to find the node correspondence between species’ protein-protein interaction (PPI) networks, has been a popular strategy to uncover missing annotations by transferring functional knowledge across species. Traditional NA methods assumed that topologically similar proteins in PPIs are functionally similar. However, it was recently reported that functionally unrelated proteins can be as topologically similar as functionally related pairs, and a new data-driven or supervised NA paradigm has been proposed, which uses protein function data to discern which topological features correspond to functional relatedness.

**Results:** Here, we propose **GraNA**, a deep learning framework for the supervised NA paradigm for the pairwise network alignment problem. Employing graph neural networks, GraNA utilizes within-network interactions and across-network anchor links for learning protein representations and predicting functional correspondence between across-species proteins. A major strength of GraNA is its flexibility to integrate multi-faceted non-functional relationship data, such as sequence similarity and ortholog relationships, as anchor links to guide the mapping of functionally related proteins across species. Evaluating GraNA on a benchmark dataset composed of several NA tasks between different pairs of species, we observed that GraNA accurately predicted the functional relatedness of proteins and robustly transferred functional annotations across species, outperforming a number of existing NA methods. When applied to a case study on a humanized yeast network, GraNA also successfully discovered functionally replaceable human-yeast protein pairs that were documented in previous studies.

**Availability:** The code of GraNA is available at https://github.com/luo-group/GraNA.

## 1 Introduction

In biomedical research, it is often challenging or infeasible to directly perform experiments on humans due to technical or ethical reasons (O’Neil *et al*., 2017). Model organisms thus have been indispensable tools for studying fundamental questions in human disease and clinical applications. Compared to humans, model organisms are simpler biological systems for comprehensive function characterization, have faster generation cycles that facilitate genetic screens, and can be readily manipulated genetically (Irion and Nüsslein-Volhard, 2022). The characterization and understanding of model organisms can provide great opportunities for translational studies in biomedicine. For example, baker’s yeast (*S. cerevisiae*) has been used as the model organism to map molecular pathways of Parkinson’s disease in humans (Khurana *et al*., 2017).

A pivotal challenge to fully realizing the potential of model organism studies for studying biomedicine is transferring the functional knowledge we learned in one species to better understand the functions of the proteins from different species (Park *et al*., 2013). A popular strategy to find functionally similar proteins is through sequence similarity search (e.g., by BLAST (Altschul *et al*., 1990)), yet sequence-similar proteins may perform different functions. In fact, it has been found that 42% of human-yeast orthologs are not functionally related (Balakrishnan *et al*., 2012; Gu and Milenković, 2021), i.e., not sharing common functional annotations. Moreover, proteins perform functions by interacting with other proteins, which form biological pathways and protein-protein interaction (PPI) networks. Therefore, in many species, similar functions can be carried out by proteins that do not have the most similar sequences but instead have similar functional roles in a biological pathway (Park *et al*., 2013). For this reason, network alignment (NA) has emerged as a complementary solution to sequence alignment for identifying the functional correspondence of proteins of different species.

Traditionally, NA aims to find the node mapping between compared networks that can reveal topologically similar regions, rather than just similar sequences. This problem is closely related to the subgraph isomorphism problem of determining whether a network is a subgraph of the other (Ullmann, 1976), which is known as NP-hard (Cook, 1971). NA of biological networks has been widely studied in bioinformatics and a large number of NA methods have been developed. Those methods circumvented the intractable complexity of the isomorphism problem by heuristically defining topological similarity based on a node’s neighborhood structure. Examples include search algorithms (Patro and Kingsford, 2012; Mamano and Hayes, 2017), genetic algorithms (Saraph and Milenković, 2014; Vijayan and Milenković, 2017), random walk-based methods (Singh *et al*., 2008; Kalecky and Cho, 2018), graphlet-based methods (Malod-Dognin and Pržulj, 2015; Milenković *et al*., 2010), latent embedding methods (Fan *et al*., 2019; Li *et al*., 2022), and many others (Meng *et al*., 2016). Moreover, in the context of the NA of social networks, graph representation learning-based methods have been proposed (Chen *et al*., 2020).

Although based on various heuristics, most existing NA methods for biological networks have a common key assumption: proteins that are in similar topological positions with respect to other proteins in the PPI network tend to have the same functions. However, observations in recent studies questioned this assumption, in which nodes aligned by those methods, while having high topological similarity, did not correspond to proteins that perform the same functions, and the topological similarity of functionally related nodes was barely higher than that of functionally unrelated pairs (Elmsallati *et al*., 2015; Meng *et al*., 2016; Guzzi and Milenković, 2018). The major reason for the failure of the assumption stems from the intrinsic noisy and incomplete nature of biological networks which contain a copious amount of spurious and missing edges. Even if we could obtain error-free PPI networks, the similar topology of cross-species subnetworks that share similar functions can be altered during evolution due to events such as gene duplication, deletion, and mutation. Therefore, solely relying on topological similarity to align biological networks may result in unsatisfactory accuracy.

Recently, Gu and Milenković (Gu and Milenković, 2020, 2021) proposed a new paradigm called data-driven NA to address the limitation of traditional NA methods. Essentially, this new paradigm transforms NA from an unsupervised problem to a supervised task, and supervised models are trained on *both* PPI network and protein function data to learn to align functionally similar nodes. The key insight is that, using function data as supervision, the model will be driven to tease topological features that are more informative for NA (termed as topological relatedness in (Gu and Milenković, 2020)) apart from other signals, such as network noise or incompleteness that are likely to break the common assumption of traditional NA methods. In contrast, most traditional NA methods are unsupervised and may not easily capture such topological features. Gu et al. have developed supervised NA methods TARA and TARA++ (Gu and Milenković, 2020, 2021), which first built graphlet features (Milenković and Pržulj, 2008) of network nodes and trained a logistic classifier with function data to distinguish between functionally related and unrelated node pairs. While outperforming traditional unsupervised NA methods, TARA (or TARA++) still has several limitations. First, its prediction performance is suboptimal as the linear logistic classifier may not be able to capture high-order, nonlinear topological features. In addition, TARA(++) is a two-stage method, where protein representations are learned in the first stage using unsupervised algorithms such as graphlet (Milenković *et al*., 2010) or node2vec (Grover and Leskovec, 2016), and in the second stage network alignment based on the learned representations is performed using supervised logistic regression. The two-stage approach may result in suboptimal alignment quality as the representation learning in the first stage is not optimized toward maximizing the alignment accuracy. Moreover, TARA(++) is not readily extended from pairwise NA to other NA problems, such as heterogeneous NA and temporal NA, for large-scale networks due to the high computational cost of counting heterogeneous or temporal graphlets (Gu *et al*., 2018; Vijayan and Milenković, 2018).

In this work, we develop GraNA, a more powerful and flexible supervised NA model for the data-driven NA paradigm for the pairwise, many-to-many network alignment problem (Guzzi and Milenković, 2018). GraNA is a graph neural network (GNN) that learns informative representations for protein nodes and predicts the functionally related node pairs across networks in an end-to-end fashion. Following TARA-TS (Gu and Milenković, 2021), GraNA also represents the two PPI networks to be aligned as a joint graph and integrates heterogeneous information as anchor links to guide the network alignment. As protein orthologs, defined as proteins/genes in different species that originated from the same ancestor, tend to retain function over evolution, GraNA further integrates across-network orthologous relationships as anchor edges to guide the alignment. One strength of GraNA is that heterogeneous data can be readily incorporated as additional nodes, edges, or features to facilitate network alignment. For example, GraNA integrates sequence similarity edges as additional anchor links to guide the alignment and pre-computed network embeddings as node features to better encode the topological roles of network nodes. GraNA is trained as a link prediction model, where function data (i.e., whether a given pair of proteins have functions in common) is used as training data. We also proposed a negative sampling strategy to improve the model training effectiveness. Since multi-modal data are integrated, GraNA is able to learn informative protein representations that reflect orthologous relationships, topology, and sequence similarity to better characterize functional similarity between proteins. Evaluated on NA tasks between five species, GraNA accurately aligned across-species protein pairs that are functionally similar. We further showed that the alignments produced by GraNA can be used to achieve accurate across-species protein function annotations. Moreover, we demonstrated GraNA’s applicability by applying it to predict the functional replacement of essential yeast genes by their human orthologs, in which GraNA re-discovered previously validated replaceable pairs in important pathways.

## 2 Methods

### Problem formulation

In this paper, we focus on pairwise NA of two species’ PPI networks. We are given as input an integrated graph *G* = (*G*_1_, *G*_2_, *E*_12_), where the undirected graph *G*_*k*_ = (*V*_*k*_, *E*_*k*_) is the PPI network of species *k* (*k* = 1, 2), with *V*_*k*_ as the set of the proteins and *E*_*k*_ as the set of physical interactions between proteins; *E*_12_ ⊆ *V*_1_ × *V*_2_ is a set of across-network edges that serve as anchor links for aligning the PPIs, such as orthologous proteins pairs. In the data-driven NA framework, the NA problem is formulated as a supervised link prediction task, where a set of functionally related protein pairs *R* = {(*u*_1_, *u*_2_)|*u*_1_ ∈ *V*_1_, *u*_2_ ∈ *V*_2_} is given as training data to train a model to predict whether a new pair of proteins are functionally related. Following previous work (Gu and Milenković, 2021; Li *et al*., 2022), the functional relatedness of two proteins is defined based on whether they share the same Gene Ontology (GO) terms (Section 3.1).

### Overview of GraNA

We propose, GraNA, a novel framework based on graph neural networks (GNNs) for supervised network alignment (Fig. 1). Receiving PPIs *G*_1_, *G*_2_, and anchor links *E*_12_ as input, GraNA first builds positional and distance embeddings as node features for every node. It then uses a GNN, which performs both within- and across-network message passing through PPI edges and anchor links (orthologs and sequence similarity), to enhance and refine those features into final representations that capture topological and evolutionary similarity relationships. GraNA is trained with protein function data to predict whether a pair of across-network proteins share the same function.

**Fig. 1.**
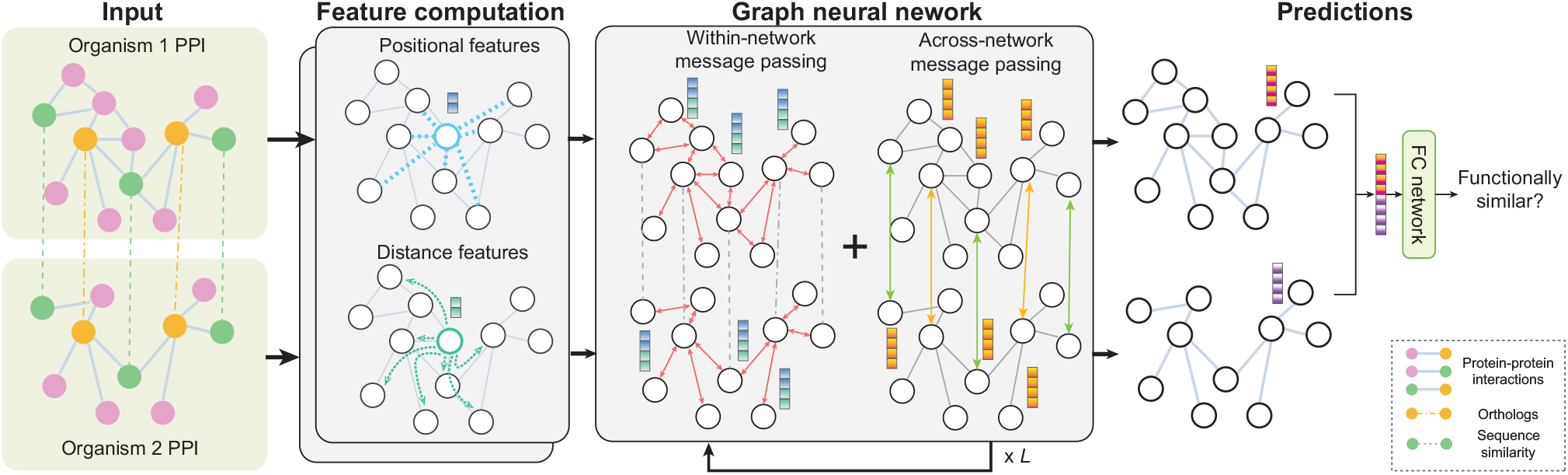
Schematic overview of GraNA. GraNA is a supervised graph neural network (GNN) that aligns functionally similar proteins in the protein-protein interaction (PPI) networks of two species. It integrates orthologs and sequence similarity relationships as anchor links to guide alignments. GraNA derives positional and distance embeddings as node features for proteins and performs iterative within- and across-network message passing to learn protein representations that capture protein functional similarity. The concatenated representations of a pair of proteins are used to make the final prediction using a fully-connected (FC) neural network.

#### 2.1 Graph neural network architecture of GraNA

GNNs have been widely used to model graph-structured data such as social networks, physical systems, or chemical molecules (Dwivedi *et al*., 2020). Here, we develop a novel GNN architecture, adapted from the Generalized Aggregation Network (Li *et al*., 2020a), to model our input PPI networks *G* = (*G*_1_ = (*V*_1_, *E*_1_), *G*_2_ = (*V*_2_, *E*_2_), *E*_12_). The key of GNNs is the graph convolution (also known as message passing) where a node first aggregates the features from its neighbor nodes, updates them with neural network layers, and then sends out the updated features to its neighbors. Through iterative graph convolutions on the PPI networks *G*_1_, *G*_2_ and anchor links *E*_12_, our model can learn an embedding for each node that encodes information of both graph topology and relationships of anchor links. GraNA has *L* layers of graph convolution blocks, where the *ℓ*-th block contains a series of non-linear neural network layers that transform node *i*’s embedding 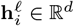 to 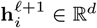, where *ℓ* ∈ [*L*] and *i* ∈ [*n*], *n* = |*V*_1_|+|*V*_2_|. In particular, h^0^ is the initialized node feature (described in Section 2.2).

Within each graph convolution block are within- and across-network propagation layers that update the node embeddings. In the *ℓ*-th block, the node embeddings 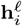 are first updated by the propagation along PPI edges (within-network message passing), in which a node aggregates its neighbor’s features using the attention mechanism:

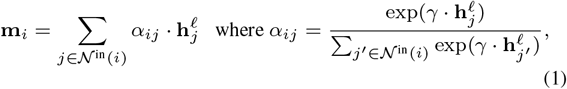

where 𝒩^in^(*i*) is the set of neighboring nodes of node *i* in terms of the within-network edge set *E*_1_ ∪*E*_2_, and *γ* is a learnable parameter known as the inverse temperature. Next, node embeddings are updated with across-network message passing through anchor links:

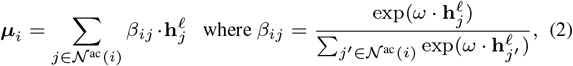

where 𝒩^ac^(*i*) is the set of neighboring nodes of node *i* in terms of the across-network edge set *E*_12_, and *ω* is a learnable temperature parameter. The final updated embedding 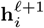 is obtained using multi-layer perceptron (MLP) *f, f*^′^ followed by a residual connection (He *et al*., 2016): 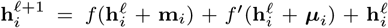. In GraNA, we stacked seven graph convolution blocks to build the GNN, where each block performs one iteration of within-network propagation (Eq. 1) and one iteration of across-network propagation (Eq. 2). Pair normalization (Zhao and Akoglu, 2019) and ReLU non-linear transformation (Nair and Hinton, 2010) are applied between two adjacent convolution blocks.

After the graph convolution, the representations of nodes *i* ∈ *V*_1_ and *j* ∈ *V*_2_, 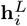 and 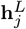, are concatenated and passed to a two-layer MLP to out a probability score that predicts whether the two nodes should be aligned. GraNA is trained using the binary cross entropy loss. The hyperparameters for training GraNA were selected based on GraNA’s performances on valid sets, and we further tested the effect of different hyperparameters on GraNA’s performances. The details of how we select our hyperparameters, the effects different hyperparameters have upon GraNA, and other implementation details are provided in the Supplementary Information.

#### 2.2 Network features of proteins

While GNNs are able to learn node embeddings that encode topological information of the input PPI network structure, previous studies have found that GNNs might perform poorly when the graph exhibit symmetries in local structure, such as node or edge isomorphism. This is related to the theoretic limitation of GNNs due to their equivalence to the 1-Weisfeiler-Lehman test of graph isomorphism (Xu *et al*., 2018). Some existing NA methods also suffered from this limitation. For example, the state-of-the-art NA method ETNA (Li *et al*., 2022) has to filter out nodes with the same neighborhood structure, since these are indistinguishable to their model when only topological information is used.

Inspired by several solutions in graph machine learning (Dwivedi *et al*., 2020; Li *et al*., 2020b), we introduce two types of node features, as the initializations of node embeddings 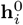, to improve the expressiveness of our GNN model and facilitate the topological feature learning. We use two complementary network features, namely the graph Laplacian positional embeddings, which encode a node’s *position* with respect to other nodes in the network, and the diffusion-based embeddings, which capture a node’s *distance* to other nodes in random walks. Intuitively, the two types of embeddings capture the long-range relationships between network nodes. In GraNA, these features are incorporated as the initialization of node embeddings 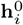 and then refined by message passing around each node’s direct neighbor vicinity. Therefore, GraNA can capture both local and global topological proximity in the network. Next, we describe how to construct the positional and distance features.

### Distance embeddings

Random walk or PageRank-based algorithms have been widely used to learn network embeddings (Perozzi *et al*., 2014; Grover and Leskovec, 2016) and improve expressiveness of GNNs (Li *et al*., 2020b). For example, the distance matrix at the equilibrium states of a random walk with restart has been used to encode the topological roles of genes or proteins in molecular networks (Cowen *et al*., 2017; Cho *et al*., 2016). Following those ideas, in this work, we compute distance embeddings for network nodes using NetMF (Qiu *et al*., 2018), a unified framework that generalizes several previous network embedding methods (Perozzi *et al*., 2014; Grover and Leskovec, 2016) and estimates the distance similarity matrix *M* in a closed form:

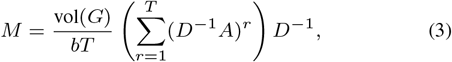

where *A* is the *n* × *n* adjacency matrix of the network, *D* is the diagonal degree matrix, 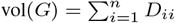 is the volume of the graph *G, b* is the parameter for negative sampling, *T* is the context window size. Unlike the adjacency matrix *A* that only contains direct neighbor relationships, the NetMF matrix *M* encodes the similarity between long-distance neighbors. The entry *M*_*ij*_ approximates the number of paths with length up to *T* between nodes *i* and *j*. In GraNA, setting *b* = 1 and *T* = 10 following the default choices (Qiu *et al*., 2018), we computed matrices *M* for the two input PPIs separately and used the *i*-th row of log *M* as the distance embedding for node *i*. As the row vector has a high dimension as the number of nodes, we applied a linear neural network layer to project the row vector from dimension *n* to *d* (*d* ≪ *n*), where *d* is the hidden dimension in GraNA’s graph convolution layers.

### Positional embeddings

In addition to distance embeddings, we further build positional features such that nodes nearby in the network have similar embeddings while distant nodes have different embeddings. For this purpose, GraNA applies the Laplacian positional encoding, which has been shown to be able to encode graph positional features in GNNs (Dwivedi *et al*., 2020). The idea is to use graph Laplacian eigenvectors that embed the graph into Euclidean space while preserving the global graph structure. Mathematically, the normalized graph Laplacian is factorized as *L* = I − *D*^−1*/*2^*AD*^−1*/*2^ = *U* ^⊤^Λ*U*, where Λ and *U* refers to the eigenvalues and Laplacian eigenvectors, respectively. In GraNA, the *d*-smallest non-trivial eigenvectors are used as the positional embeddings and concatenated with the distance embeddings together as the initialized node features 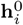.

#### 2.3 Integrating heterogeneous information for NA

In addition to PPIs, there are other types of relationships that can help characterize the functional similarity of proteins, such as gene-gene interactions, sequence similarity, phenotype similarity, and associations between proteins and other entities such as diseases. A naive way to integrate multiple data sources is to collapse them as additional but the same type of nodes and edges in a flattened network, which, however, may lose context-specific information. Heterogeneous data integration, which treats distinct types of nodes and edges separately, has been shown effective to integrate diverse data sources (Cho *et al*., 2016; Luo *et al*., 2017). A few previous NA studies consider the heterogeneous NA problem, but their approaches required non-trivial modifications in the optimization objective and feature engineering as compared to the homogeneous NA problem. On the contrary, one of the major advantages of GraNA is that it can readily integrate heterogeneous information to facilitate the alignment of networks by simply including the data as additional nodes, edges, or feature embeddings and applying heterogeneous graph convolutions to capture context-specific information.

As a proof-of-concept, here we apply GraNA to incorporate sequence similarity relationships as another type of anchor links in addition to the orthologous relationships. Now we have two sets of across-network edges as input, which are denoted as 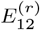 for *r* = 1, 2. To learn embeddings from heterogeneous data, we perform separate across-network message passing for each edge type: 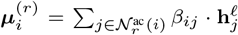. Compared to Eq. 2, note that the aggregation and the weights *β*_*ij*_ here are defined on *i*’s neighbor nodes that are connected by the *r*-th type of edges 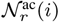, instead of all neighbors 𝒩(*i*). After performing both types of message passing, we obtain the updated node embedding using a sum pooling operation over all edge types: 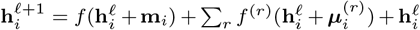, where *f* ^(*r*)^ is a fully connected neural network specific to edge type *r*. We expect that, by multi-view information from orthologs and sequence similarity edges, GraNA can better distill the topological features that are useful to predict functional relatedness. Of note, GraNA is a generic framework, and other types of node or edge data can be integrated into GraNA in a similar way.

#### 2.4 Enhancing model learning with hard negative sampling

Supervised NA essentially is a positive-unlabeled learning problem, meaning that we only observed positive protein pairs that are functionally related (e.g., have at least one GO term in common), denoted as ℐ_*p*_ = {(*p, q*)|proteins *p* and *q* are functionally related}, without observing validated negative samples. For a new pair (*p*^*^, *q*^*^) ∉ ℐ_*p*_, it does not necessarily mean that the two proteins do not have the same function, rather, it is more likely their functions have not been thoroughly characterized by experiments. To generate negative samples for training a supervised classifier to distinguish functionally related and unrelated pairs, previous NA methods usually chose to randomly sample a set of pairs not in ℐ_*p*_ as the negative set ℐ_*n*_ (Gu and Milenković, 2021, 2020).

We reason that the random negative sampling might lead the machine learning model to learn the node’s presence in the training data rather than functional relatedness. Denote 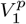 as the set of proteins in PPI *G*_1_ that are involved in positive pairs, i.e., 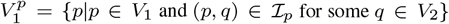, and 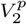 has a similar meaning. As only a small fraction of proteins in *V*_1_ and *V*_2_ are involved in the positive set, for most randomly-sampled negative pairs (*p, q*), it is likely that *p* and *q* are new proteins that did not occur in 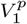 or 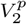, respectively. Due to the distribution discrepancy between positive and negative samples in the training set, machine learning models trained on this data may only learn to predict for a given pair of proteins (*p, q*) whether 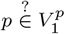 or 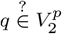, rather than predicting 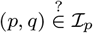.

To encourage the model to learn the functional relatedness instead of node representativeness in the training data, we propose a hard negative sampling strategy to construct the negative set, where the sampled negative edges must contain nodes that have both appeared in positive edges. We achieve this by performing edge swap between positive edges: given two positive pairs (*p*_1_, *q*_1_) and (*p*_2_, *q*_2_), we swap their endpoints and add new edges (*p*_1_, *q*_2_) and (*p*_2_, *q*_1_) to the negative set if there did not show in ℐ_*p*_. Equivalently, the set of negative edges is defined as 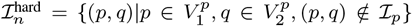. In our experiments, we also compared to two other negative sampling strategies, including the “easy” sampling used in previous NA studies (Gu and Milenković, 2020, 2021): 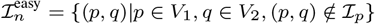, and a “semi-hard” sampling that requires a sampled negative edge to contain at least one node that has appeared in positive edges : 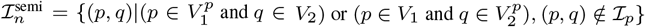.

## 3 Results

We performed several experiments to assess GraNA’s ability to capture the functional similarity of proteins and predict protein functions across species. We also conducted ablation studies to better understand the model’s prediction performance. Furthermore, we used a proof-of-concept case study to demonstrate GraNA’s applicability in functional genomics.

### 3.1 Datasets

#### Network data

The PPI network data of six species (*Saccharomyces cerevisiae, Schizosaccharomyces pombe, Homo sapiens, Caenorhabditis elegans, Mus musculus*, and *Drosophila melanogaster*) were downloaded from BioGRID (v3.5.187) (Stark *et al*., 2006). We used both orthologs and sequence similarity relationships as anchor links to guide network alignment. For orthologs, we followed the ETNA study (Li *et al*., 2022) and downloaded orthology data from OrthoMCL (v6.1) (Li *et al*., 2003). For sequence-similar pairs, we retrieved the expert-reviewed sequences, if any, of proteins in our PPI networks from the UniProtKB/Swiss-Prot database (Consortium, 2023). We then used MMseqs2 (Steinegger and Söding, 2017) to perform sequence similarity searches between the proteins of pairwise species and kept protein pairs with an E-value ≤ 10^−7^ as anchor links. We chose this cutoff following previous work (Gu and Milenković, 2021; Kalecky and Cho, 2018), and we observed that varying this cutoff in a wide range had no significant impact on our GraNA’s prediction performance (Fig. S1). The statistics of the PPI networks and the anchor links can be found in Supplementary Tables S1 and S2.

#### Functional annotations

We collected the functional annotations (terms) from Gene Ontology (GO) (Ashburner *et al*., 2000) (2020-07-16) and considered two proteins to be functionally similar if their corresponding genes have the same GO terms. Following ETNA (Li *et al*., 2022), we only kept annotations related to the Biological Process (BP) category, which are propagated through *is a* and *part of* relations, and included evidence codes EXP, IDA, IMP, IGI, and IEP. As GO terms appearing at the higher levels of the GO hierarchy might be too general or redundant, following ETNA (Li *et al*., 2022) and other studies (Gu and Milenković, 2020, 2021), we focused our analyses on specific functions by creating a slim set of GO terms associated with at least 10 genes but no more than 100 genes. Another expert-curated GO slim terms were also added to this slim set (Greene *et al*., 2015). The statistics of functionally similar protein pairs between species can be found in Supplementary Table S2.

### 3.2 GraNA better exploits topological similarity for network alignment

We first assessed GraNA’s ability for network alignment by applying it to align the networks between human and four major model organisms, including *S. cerevisiae, M. Musculus, C. elegans*, and *D. melanogaster*, and between two yeast species (*S. cerevisiae* and *S. pombe*). The prediction task was formulated as a link prediction problem, i.e., predicting whether two proteins have the same function. We created an out-of-distribution train/test split (with a ratio of 8:2) such that proteins present in the training set never occur in the test set. In another more challenging split, we further forced that the training proteins and test proteins do not have *>* 30% sequence identity.

We compared GraNA to two unsupervised embedding-based methods (ETNA (Li *et al*., 2022), MUNK (Lim *et al*., 2018)), a graph theoretic method (IsoRank (Singh *et al*., 2008)), a sequence similarity-based method (MMseqs2 (Steinegger and Söding, 2017)), and two supervised methods (TARA-TS and TARA++ (Gu and Milenković, 2021)). We used the same PPI networks and orthologs anchor links for all baseline methods. Anchor links for protein pairs that share GO terms were removed to avoid data leakage. To make a fair comparison, we included a variant of our method (GraNA-o) that only used orthologs (without sequence-similar pairs) as anchor links. Unsupervised methods were evaluated on the same test set used for supervised methods. The running time analyses of GraNA and representative baseline methods can be found in Supplementary Table S6. The evaluation results suggested that GraNA consistently outperformed other methods for aligning functionally related proteins in all five NA tasks in terms of the AUROC and AUPRC metrics (Fig. 2). Precision, recall, and total number of predicted alignments were reported in Figs. S11, S12, and S13. We first confirmed the advantage of the supervised NA paradigm over the traditional unsupervised paradigm: GraNA(-o) substantially improved other unsupervised methods (MMseqs2, IsoRank, MUNK, and ETNA) with clear margins. For example, the AUROC and AUPRC improvements achieved by GraNA over the best unsupervised method (ETNA) were 11% and 55%, respectively (averaged over five tasks). Compared to those unsupervised methods that entirely rely on the topology to align nodes and are susceptible to the noise and incompleteness in biological networks, GraNA further leveraged function data as direct supervision signals to tease topological features that are directly related functional relatedness from background noise and greatly improved the alignment quality.

**Fig. 2.**
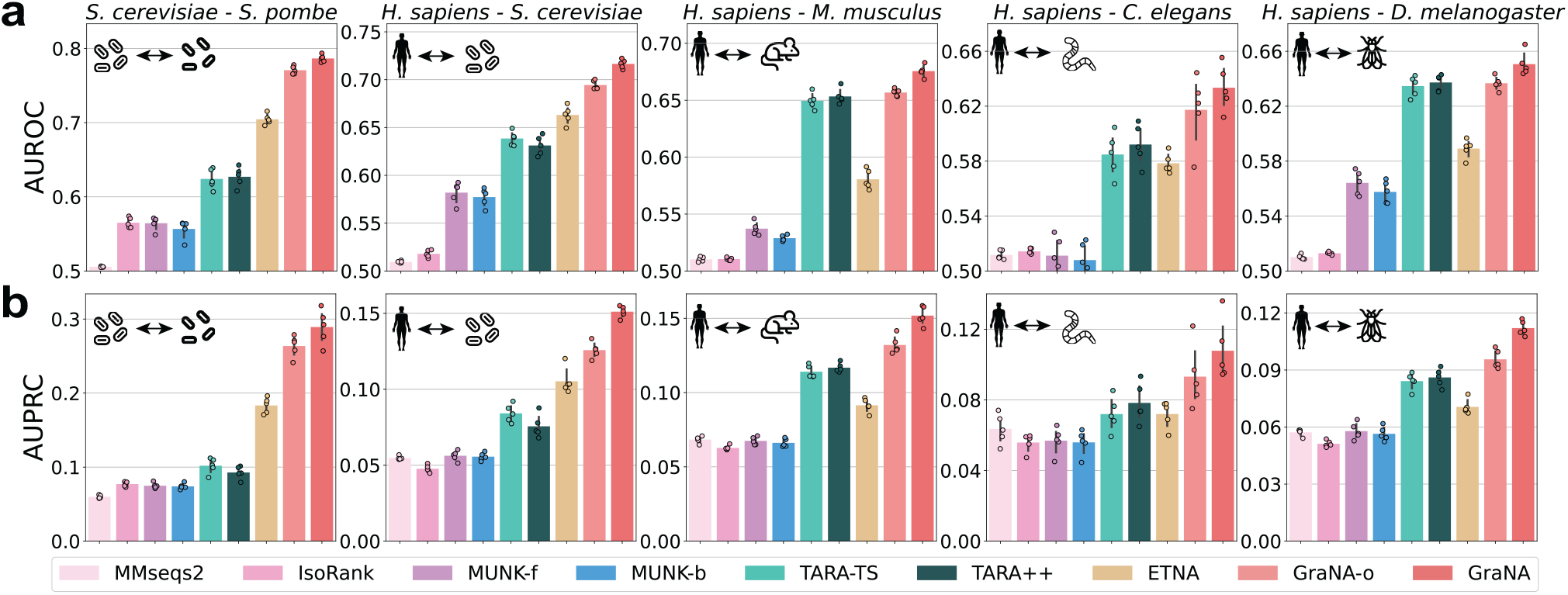
Performances of network alignment prediction. GraNA and other baselines were evaluated for aligning functionally similar proteins across five pairs of species, using (a) AUROC and (b) AUPRC as metrics. GraNA-o is a variant of GraNA that only uses orthologs as anchor links whereas GraNA refers to the full model that uses both orthologs and sequence similarity as anchor links. The default E-value cutoff 10-3 is used for MMseqs2. As MUNK is not a bidirectional NA method, the performances of its forward and backward predictions were shown separately as MUNK-f and MUNK-b. Performances were evaluated using five independent train/test data splits. Raw AUROC and AUPRC scores are provided in Tables S4 and S5.

In addition, compared to TARA-TS and TARA++, the only methods for the supervised NA paradigm in literature, we found that our method is a more powerful deep learning solution for supervised NA. For example, GraNA-o on average had 53% higher AUPRC scores than TARA-TS. Interestingly, TARA-TS, despite as a supervised method, sometimes even had a lower performance than the state-of-the-art unsupervised method ETNA. The potential reason is that TARA-TS only used a linear logistic model that only able to model linear feature interactions in the data, while GraNA is an end-to-end graph neural network, which captures more complex, non-linear feature dependencies, and can better exploit topological similarity and predict node alignment.

Moreover, although GraNA-o outperformed other methods in most scenarios, in a few cases, it was only on par with the second-best baseline (TARA-TS; Fig. 2, 3rd and 5th columns). However, we found that when integrating both orthologs and sequence similarity as anchor links, the full model (GraNA) further improved GraNA-o and outperformed all other baselines in all tasks in both AUROC and AUPRC, suggesting that GraNA was an effective tool to integrate heterogeneous data for boosting the network alignment performance. In contrast, we observed that TARA-TS, even when given the two types of anchor links, was not able to improve the alignment performance (to be discussed in Section 3.4 and Fig. 4a).

On a more challenging data split where the sequences in the train and test sets have no sequence identity *>* 30%, we also observed that GraNA clearly outperformed the second best baselines ETNA and TARA-TS (Fig. S3 and S4). We also had similar observations when using other sequence identity cutoffs to create the train/test splits (Fig. S2). This strict benchmark suggested that GraNA can generalize its prediction for proteins that are sequence-dissimilar from what it has seen in the training data.

Additionally, we created another challenging evaluation dataset based on a temporary split strategy, where the snapshot of the GO database as of 2018-07-02 was used as training data, and the GO snapshot as of 2022-12-04, excluding all training annotations, was used as test data. On this dataset, we again observed similar results where GraNA outperformed baselines such as ETNA and TARA-TS (Fig. S5). This demonstrated GraNA’s generalizability when making predictions for proteins whose functions are not completely characterized.

Overall, these results demonstrated that GraNA can better explore topological similarity to accurately align networks. The flexible GNN framework further allowed GraNA to integrate heterogeneous data types that capture multi-view similarity relationships to improve the alignment quality.

### 3.3 GraNA translates accurate network alignments to function predictions

One important application of NA is to better understand human protein functions by transferring our learned function knowledge about model organisms. Therefore, after evaluating the performance of aligning functionally related proteins, we next studied whether the network alignments produced by GraNA can facilitate protein function prediction. Here, we applied GraNA to generate the alignments between humans and the four model organisms. Then, we considered the top 5,000 ranked protein pairs aligned by GraNA and calculate the Jaccard index between the functional annotations of the two proteins in each pair. As a protein may have multiple functions, this evaluation aimed to quantify the overlap between the sets of functions of the two aligned proteins, which was more complex and challenging than the evaluation in the last section which predicted whether two proteins share at least one function. We also compared a random baseline that randomly samples 5,000 pairs from proteins that have at least one GO term, in addition to our previously introduced baselines. Furthermore, we have also evaluated GraNA in an established protein function prediction framework (Meng *et al*., 2016).

We observed from Fig. 3 that, even with the partial model GraNA-o, our method has already outperformed other methods on three out of the four tasks in terms of Jaccard similarity. The full model GraNA, which integrated heterogeneous orthologs and sequence similarity edges, further boosted the function prediction performance. These results suggested that GraNA was able to not only align functionally similar protein pairs but also prioritize “most similar” pairs to the top of its prediction list. GraNA’s ability to prioritize functionally similar proteins has important implications when studying human diseases, since it can suggest the most functionally similar counterpart of a human gene in model organisms for detailed characterization. Moreover, we noted that the improvements achieved by GraNA over other methods were more pronounceable for species with high-quality PPI networks (e.g., *S. cerevisiae*). On the alignment task between human (*H. sapiens*) and roundworm (*C. elegans*), GraNA achieved performance on par with the second best baseline, which was likely due to that the PPI of *C. elegans* is the sparsest among all four model organisms (density *<* 0.2%). This finding was consistent with the ETNA study (Li *et al*., 2022). We also compared GraNA with other methods using the functional coherence (FC) metric (Singh *et al*., 2008; Chindelevitch *et al*., 2013), a variant of the Jaccard index that only focuses on standardized GO terms to avoid bias caused by terms from different levels of the GO hierarchy, and observed similar performance (Fig. S10). Additionally, by using the protein function prediction framework (Meng *et al*., 2016), we observed that GraNA predicted a smaller set of predictions with higher precision compared to TARA++, which is useful when high-confidence and limited false positive predictions are desired (Fig. S14). Overall, this experiment here suggested that GraNA translated its effective network alignments to the accurate predictions of protein functions, demonstrating its potential for across-species functional annotations.

**Fig. 3.**
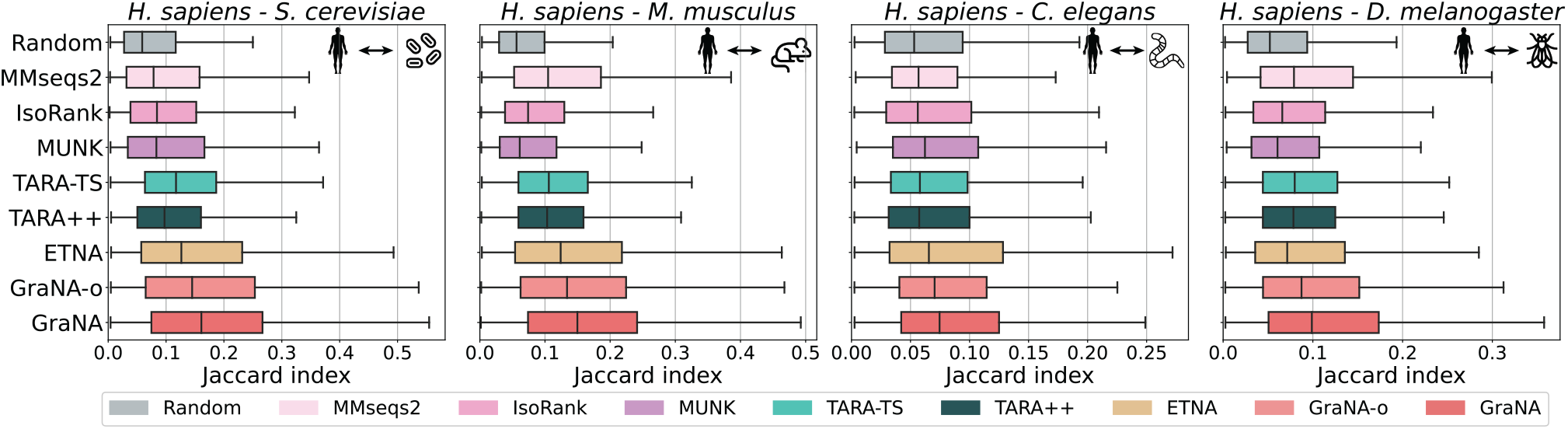
Performance of protein function prediction. Based on the network alignments produced by each method for four pairs of species (*H. sapiens*-*S. cerevisiae, H. sapiens*-*M. Musculus, H. sapiens*-*C. elegans, H. sapiens*-*D. melanogaster*), we chose the top 5,000 ranked protein pairs and transferred all the functional annotations of one protein in an aligned pair to predict the other protein’s function. The accuracy of the function prediction was evaluated by calculating the Jaccard index between the sets of the two aligned proteins. Box plots showed the distribution of the Jaccard index of the top 5,000 aligned pairs for each method on five NA tasks.

### 3.4 Analyses of key model designs in GraNA

Having validated that GraNA outperformed state-of-the-art methods for aligning networks and predicting functions, we performed ablation studies to understand the GraNA model in more detail and attribute performance improvements to several key design choices in GraNA.

#### Heterogeneous anchors

As GraNA is a flexible framework to integrate heterogeneous data, we first investigated the effects of using heterogeneous data on the performance of network alignment. We compared GraNA variants that used only orthologs, only sequence similarity, or both as anchor links. We observed that with either of the anchor links, GraNA was able to achieve an AUPRC better than the two best baselines (ETNA and TARA-TS) and combining both of them led to the best AUPRC score (Figs. 4a and S6). Interestingly, we found the two baselines, when given two types of anchors, did not improve their network alignment accuracy compared to when a single type of anchor was used (Figs. 4a and S7). These comparisons indicated that information contained in the two types of edges are not redundant but complementary, and GraNA can integrate them more effectively than other baselines. The major reason was that GraNA implemented separate message passing mechanisms to handle different types of anchors, while ETNA and TARA-TS (with node2vec features (Grover and Leskovec, 2016)) mixed them as a single type of edges. We expect that integrating more data that capture multiple aspects of protein similarity can further help GraNA to better characterize protein functional relatedness.

**Fig. 4.**
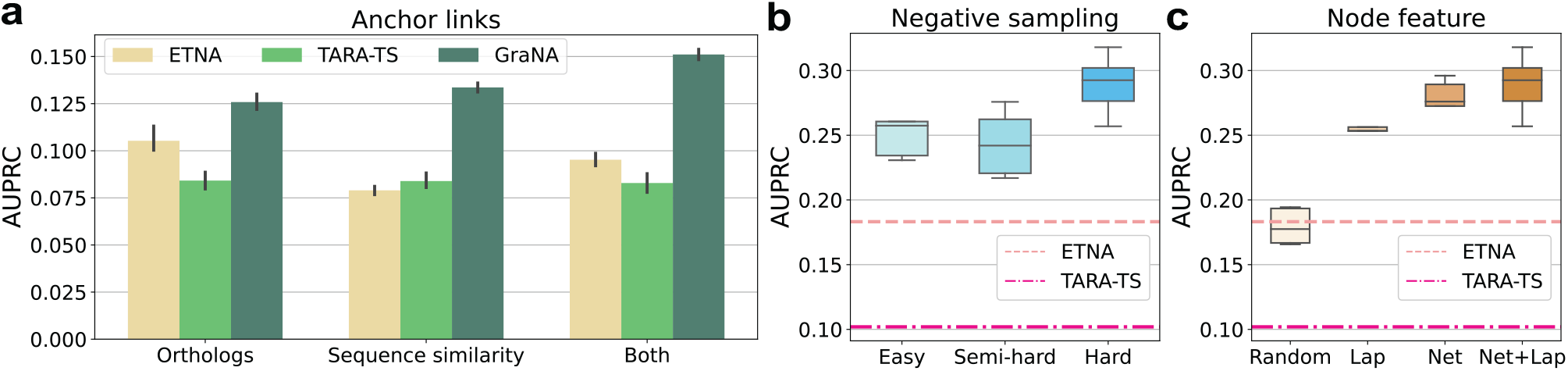
Ablation studies validated key designs of GraNA. (a) Comparison between GraNA and two of the best baselines, TARA-TS and ETNA on heterogeneous data integration, where either orthologs, sequence similarity, or both were used as the anchor links for the network alignment between *H. sapiens* and *S. cerevisiae*. (b-c) Ablation analyses that compared different negative sampling strategies (b) and node features (c) for the network alignment between *S. cerevisiae* and *S. pombe*. AUPRC scores of the two best baselines (ETNA and TARA-TS) were shown in (b) and (c) for reference. Performances were based on five independent trials of train/test split. Comparisons on all species can be found in Supplementary Information. (Lap: Laplacian embeddings; Net: NetMF embeddings.)

#### Hard negative sampling

Another novel design in GraNA is the hard negative sampling which prevented the model from only learning from trivial training samples. To better illustrate this, we compared GraNA models trained with three negative sampling strategies, including easy, semi-hard, and hard negative sampling (Methods). We observed that GraNA trained with easy and semi-hard samplings already outperformed the second-best baseline, and using the hard sampling further improved the AUPRC by 20% and showed a more significant margin over baselines (Fig. 4b). Hard negative sampling is a critical ingredient that makes GraNA accurate and generalizable. As discussed in the Methods section, random negative sampling tends to create a training set that confuses the machine learning model, and the model may just learn whether protein appeared in the training set rather than the functional relatedness between protein pairs. In contrast, hard negative sampling forces our model to discriminate between functionally related and unrelated pairs.

#### Node features

We used both distance features (NetMF embeddings) and positional features (Laplacian embeddings) to initialize the node features in GraNA. Here, we analyze the effect of the node features by comparing GraNA variants that used only one or both of the NetMF and Laplacian embeddings, or randomly initialized node features. We observed that with random node features, the prediction performance was only comparable with the unsupervised ETNA method (Fig. 4c). When replacing the random features with network-informed features (NetMF and Laplacian), GraNA significantly improved its AUPRC scores. Finally, incorporating both embeddings led to the highest AUPRC score. This comparison underscored the effectiveness of using informative features. Although the GNN model alone was able to capture topological properties of network nodes, it still only captured localized information as a node’s features were only propagated to its nearby neighbors with a few times (e.g., *<* 10) of message passing. However, the two embeddings we used were able to encode global, long-range neighbor relationships between nodes, which were complementary to the topological features learned by the GNN and jointly enhanced GraNA’s effectiveness.

### 3.5 Application: predicting replaceability for a humanized yeast network

Finally, we demonstrate the applicability of GraNA using a task of identifying replaceable human-yeast gene pairs. Recent studies have identified many human genes that can substitute for their yeast orthologs and sustain yeast growth (Kachroo *et al*., 2015; Laurent *et al*., 2020), which provides a tractable system known as ‘humanized yeast’ to allow for high-throughput assays of human gene functions. Given that not all yeast genes can be replaced by their human orthologs, biological NA methods might become useful tools to predict the replaceability among human-yeast orthologs.

We collected the experiment data from Kachroo et al. (Kachroo *et al*., 2015), which has assayed 414 essential yeast genes for complementation by their human orthologs and found 47% of them could be humanized. After filtering out genes that are not included in the PPI network of *S. cerevisiae* that we used in this work, we obtained 411 gene pairs, out of which 174 replaceable pairs are labeled as positive samples and the remaining as negative. To avoid potential signal leakage, in our data we further removed 169 orthologs that coincide with the 411 pairs. Using this data as a binary classification test set, we first applied a baseline method, ETNA, to predict the replaceability of each human-yeast pair. We observed that ETNA’s predicted performance was nearly random (AUC∼0.5; Fig. 5a). This was not surprising because, by design, ETNA was trained to classify between orthologs and non-orthologs, while all the positive and negative pairs in the test sets here are all human-yeast orthologs, which appeared to be indistinguishable to ETNA. Next, we applied the GraNA model pre-trained on our *H. sapiens*-*S. cerevisiae* alignment task (GraNA-pt) to predict for those 411 gene pairs. Even though GraNA-pt was not directly trained to predict replaceability, we found that it still had a better-than-random prediction accuracy (AUC=0.56; Fig. 5a) on the test set, which suggested that the functional similarity relationships captured by GraNA were relatively more generalizable. After fine-tuning the trained GraNA model on the 411 gene pairs by re-training the parameter of the top MLP layers and freezing GNN layers, we observed that this model (GraNA-ft) reached an AUC of 0.68 in five-fold cross-validation (Fig 5a), which was higher than the AUC of the supervised TARA-TS model (Fig S8). This suggested that the prediction accuracy of GraNA on this task could be improved with direct supervision.

**Fig. 5.**
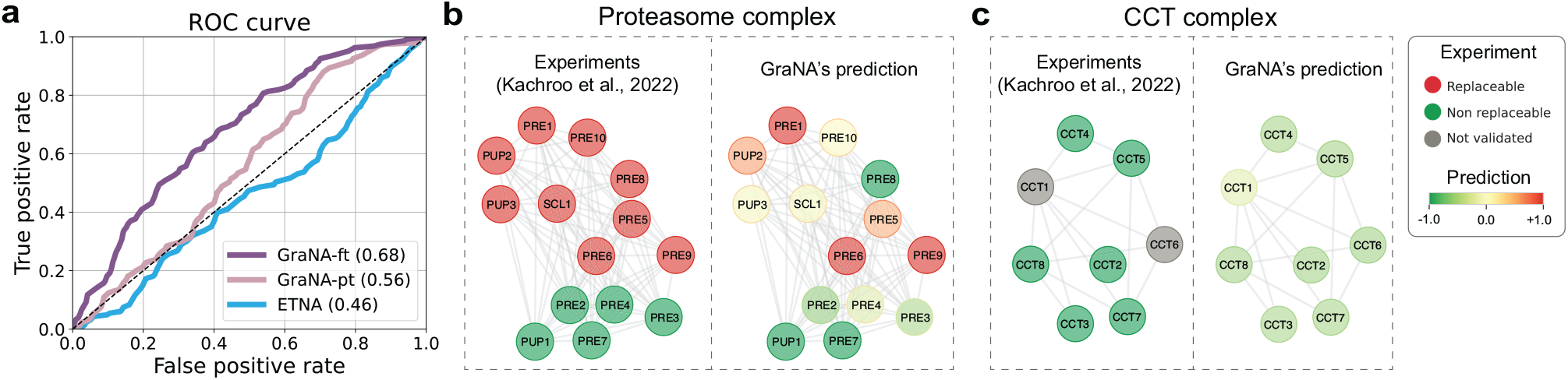
An application of GraNA on predicting replaceable human-yeast gene pairs in humanized yeast network. GraNA was used to predict whether human genes can replace their yeast orthologs for the functions in humanized yeast network. (a) Using 414 validated positive and negative human-yeast gene pairs by Kachroo et al. (Kachroo et al., 2015) as the test set, we compared a pre-trained GraNA model (GraNA-pt), a fine-tuned GraNA model (GraNA-ft), and ETNA to evaluate their ability to distinguish replaceable and non-replaceable human-gene pairs in the test set. The performance of GraNA-ft was evaluated using five-fold cross-validation whereas GraNA-pt and ETNA were evaluated on the whole test set directly. (b-c) Case studies where GraNA was used to predict the replaceability in two pathways: (b) Proteasome complex and (c) CCT complex. We visualized the complex validated by Kachroo et al. (Kachroo et al., 2022) for comparison, where validated replaceable genes were colored in red, non-replaceable genes in green, and unvalidated genes in gray. For GraNA’s predicted network, the predicted score for each gene was normalized into a z-score and colored with a gradient colormap from green (most non-replaceable) to red (most replaceable).

As a case study, we applied GraNA to study the replaceability in protein complexes. We selected as two examples the proteasome complex and the CCT complex that have experimental validation data (Kachroo *et al*., 2022, 2017, 2015). In both examples, we used the genes in the complex as the test set and the remaining genes with experimental validation data as the training set. For the Proteasome complex that contains both replaceable and non-replaceable genes, except for *PRE8*, GraNA correctly predicted a positive z-score for replaceable genes and a negative z-score for non-replaceable genes (AUC=0.91; Fig. 5b). For the CCT complex that was mainly enriched with non-replaceable genes, GraNA’s prediction also recapitulated the replaceability in the network, where validated non-replaceable genes were predicted with a negative z-score (Fig 5c).

Overall, these results demonstrated the applicability of GraNA for extending the network alignments to empower other functional analyses of genes and proteins.

## 4 Conclusion

NA is a fundamental problem in various domains, such as linking users across social network platforms (Zafarani and Liu, 2013), unifying entities across different knowledge databases (Zhu *et al*., 2017), and aligning keypoints in computer vision (Sarlin *et al*., 2020). In this paper, we studied the NA problem for biological networks. We have presented GraNA, a deep learning model for aligning functionally related proteins in cross-species PPI networks. Our work was motivated by the recently proposed supervised network alignment methods such as TARA/TARA-TS (Gu and Milenković, 2020, 2021), which represent the two PPIs being aligned as a joint graph connected by anchor links and integrate topology, sequence, and function information to characterize the function similarity between cross-species protein pairs. GraNA integrates PPI networks, ortholog and sequence similarity relationships, network distance and positional embeddings, and protein function data to learn to align across-species proteins that are functionally similar. Experiments showed that GraNA outperformed state-of-the-art NA methods, including both supervised and unsupervised approaches, on aligning pairwise PPI networks for five species, and the high-quality network alignments of GraNA also enable accurate functional prediction across species. We further investigated several key model designs of GraNA that led to performance improvements and demonstrated the applicability of GraNA using a case study of predicting replaceability in humanized yeast network. GraNA is a flexible framework and can be readily extended in the future to integrate diverse types of entity and association data to facilitate NA. As previous methods such as TARA, GraNA can also be generalized to study other NA problems, including multi-species NA and temporary NA.

## Acknowledgement

Y.L. is partially supported by a 2022 Seed Grant Program of the Molecule Maker Lab Institute and a 2022 Amazon Research Award. We thank the authors of ETNA (Li *et al*., 2022) for sharing the benchmark datasets. Species icons in Figures 2 and 3 were downloaded from https://www.flaticon.com.

## 1 Supplementary Information

### 1.1 Experimental setup

For network alignment prediction, GraNA was trained on the training set, tuned on the valid set, and evaluated on the test set along with other NA baselines, where the split ratio is 70/10/20. For each dataset, data split was performed 5 times with 5 different seeds.

In the context of NA for biological networks, many different evaluation metrics have been proposed, and they often focus on different aspects of network alignment prediction. Ma and Liao (2020) categorized some of the most commonly used metrics into two types: biological evaluation and topological evaluation. Apart from the metrics summarized by Ma and Liao (2020), Fan *et al*. (2019); Li *et al*. (2022) also used AUPRC and AUROC for evaluating the predicted network alignment. In this work, we selected metrics following Singh *et al*. (2008); Chindelevitch *et al*. (2013); Fan *et al*. (2019); Li *et al*. (2022) and included AUROC, AUPRC, Jaccard index (also known as Gene Ontology Consistency), and functional coherence (FC) as metrics. As the prediction of GraNA is a many-to-many mapping between the across-species proteins, we cannot directly leverage a particular set of metrics used in previous studies Saraph and Milenković (2014); Vijayan and Milenković (2017), such as Edge Correctness (EC) and Node Correctness (NC), which are based on the assumption that the mapping is one-to-one and require non-trivial modification for our purpose.

For protein function prediction, we chose the Jaccard index and functional coherence of the top 200 predicted node pairs given for each train/test split (1000 pairs in total after combining the top pairs from five runs). For a fair comparison, we filtered anchor links that coincide with positive test pairs from our training data. Jaccard index describes how similar two proteins are in terms of function, as it is calculated as |*S*_1_ ∩ *S*_2_ |*/*|*S*_1_ ∪ *S*_2_ |, where *S*_1_, *S*_2_ represent respectively the set of GO terms the two nodes are annotated with. Following previous studies (Singh *et al*., 2008; Chindelevitch *et al*., 2013), we define functional coherence as follows. GO terms were first mapped to a standardized GO set. Within this set, all GO terms are at a distance of 5 to the root of the GO hierarchy, and any GO terms with a distance less than 5 to the root are dropped. We measured the distance only by considering the relations *is a* and *part of* in Biological Process (BP) of the GO, and we retrieved the ancestor information of each GO term through the QuickGO REST API (Binns *et al*., 2009). This design aimed to avoid evaluating functional similarity at different levels of the Gene Ontology graph. For each protein pair (*x, y*), functional coherence is defined as |*S*_*x*_ ∩ *S*_*y*_ |*/*|*S*_*x*_ ∪ *S*_*y*_ |, whereas *S*_*x*_, *S*_*y*_ represent the sets of standardized GO terms with protein *x, y* respectively. Using Jaccard index and functional coherence, we are able to quantify the proportion of functional knowledge that is successfully transferred from the network alignment established by GraNA.

### 1.2 Baselines

In experiments, we compare GraNA with several existing NA methods. For a fair comparison, all baseline methods were trained, if needed, and evaluated on the same data as GraNA. Specifically, for baselines that require anchor links, we used the same ortholog anchor links that GraNA uses. Default parameters were used for all baselines.

For unsupervised NA method, we included IsoRank (Singh *et al*., 2008), MMseqs2 (Steinegger and Söding, 2017), MUNK (Fan *et al*., 2019), and ETNA (Li *et al*., 2022). MMseqs2 is a tool for calculating sequence similarity and clustering proteins based on their sequences. We included it as a baseline method for assessing the relatedness of sequence similarity to functional similarity. IsoRank is an unsupervised multi-network alignment method, which is based on the intuition that functionally similar proteins have similar sequences and neighborhood topologies. The alignment of networks is formulated as an eigenvalue problem. IsoRank was originally designed to align orthologous pairs using sequence similarity as anchor links. MUNK, linking two PPIs via orthologs, uses matrix factorization to create a functional embedding in a way that proteins from different species are embedded in the same space. Then, a score matrix is calculated between two species, which can be used for network alignment prediction. ETNA is the state-of-the-art unsupervised NA method. It first learns representations for proteins from the PPIs via autoencoder and then applies a cross-training mechanism using orthologs to align the embeddings from two species. For the supervised NA method, we included TARA-TS and TARA++ (Gu and Milenković, 2021). From the three versions of TARA-TS (graphlet (Milenković and Pržulj, 2008), node2vec (Grover and Leskovec, 2016), metapath2vec (Dong *et al*., 2017)), we chose the version based on node2vec as it showed the best performance among the three as shown in their experiments. Regarding TARA++, for the protein function prediction evaluation framework (Meng *et al*., 2016), we implemented TARA++ according to its original definition, which is the intersection of TARA and TARA-TS predictions. For network alignment prediction, we had to make a tweak on TARA++: in the TARA++ paper, TARA++ was developed for the function prediction task but not for the network alignment task. Therefore, we adapted TARA++ to the network alignment prediction to compare with GraNA – we first ran TARA and TARA-TS to obtain the predicted probability (produced by the logistic regression classifier) that a given protein pair shares at least one GO term and then we took the average to TARA’s and TARA-TS’s predicted probabilities. The averaged probability was used as the prediction of TARA++. The average operation here followed the same idea of the intersection operation in the original TARA++ for function prediction, which took the consensus predictions of TARA and TARA-TS. In addition to the average, we have tried combining TARA and TARA-TS by taking their minimum or maximum predicted probability for network alignment, and the results were similar. Using this approach, we were able to compare TARA++ to other methods in our network alignment benchmark.

### 1.3 Hyperparameters

The hyperparameters in GraNA include the total number of epochs, batch size, learning rate, hidden dimension, number of graph convolution blocks, and graph convolution type. We comprehensively tested the robustness of GraNA against different hyperparameter settings. The search space of hyperparameters for training GraNA was shown in Table S3. For each train/valid/test split, GraNA was first trained on the training set and then validated on the validation set. We chose the final combination of hyperparameters for training GraNA based on GraNA’s performances (AUROC and AUPRC) on the validation set. To avoid an exponential number of combinations of hyperparameters that would make the grid search infeasible, we fixed the values of other hyperparameters when tuning one specific hyperparameter.

We evaluated four different types of graph convolution layer: GCN (Kipf and Welling, 2016), SAGE (Hamilton *et al*., 2017), GAT (Veličković *et al*., 2017), and GEN (Li *et al*., 2020a). The four architectures differ from each other mainly in their neighborhood information aggregation mechanisms. GCN aggregates neighborhood information in a weighted mean manner based on node degrees and edge weights from the normalized Laplacian matrix. SAGE, in comparison, takes a mean over neighborhood node features for constructing the message for one node. GAT employs the attention mechanism for aggregating node features, whereas GEN is the layer we used in GraNA, and it aggregates neighborhood information through a softmax function.

Raw results averaged on five independent train/valid split for each hyperparameter setting for alignment between *S. cerevisiae* and *S. pombe* were shown in Figure S9. We observed that GraNA was robust to hyperparameters. Given the results of hyperparameter tuning and the computational resources available to us, we built a total of 7 graph convolution blocks, each with a hidden dimension of 128 and a convolution type of GEN (Li *et al*., 2020a), for GraNA. During training, we used the Adam optimizer with an initial learning rate of 0.001 and a weight decay of 5e-4, and we set the batch size to be 2^16^. We trained GraNA for a maximum of 200 epochs. GraNA was trained on a single NVIDIA A40 GPU card. The running time analyses of GraNA and baseline methods (TARA, TARA-TS, ETNA) are provided in Table S6.

## 2 Supplementary Tables

**Table S1.** The number of nodes and edges in the PPI network of each species. Abbreviations: sce: *S. cerevisiae*, spo: *S. pombe*, hsa: *H. sapiens*, mmu: *M. Musculus*, cel: *C. elegans*, and dme: *D. melanogaster*.

| Type | sce | spo | hsa | mmu | cel | dme |
| --- | --- | --- | --- | --- | --- | --- |
| Nodes | 5,669 | 2,334 | 17,120 | 7,762 | 4,439 | 7711 |
| Edges | 110,776 | 10,525 | 418,512 | 47,833 | 18,301 | 49,769 |

**Table S2.**
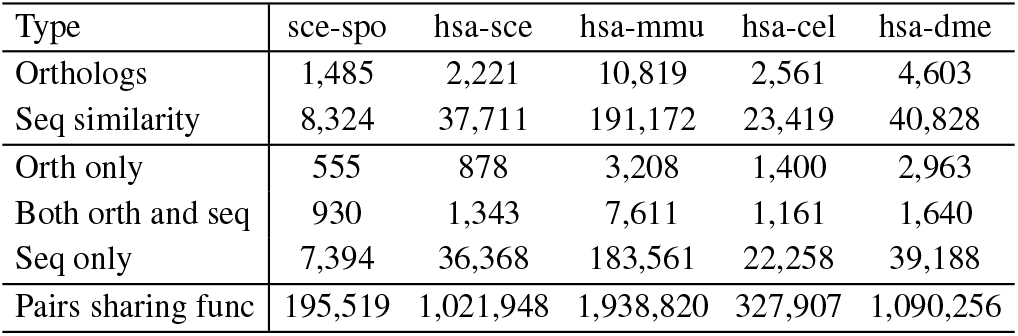
The number of anchor links (orthologs and sequence similarity) and protein pairs sharing function in each pair of PPI networks. *Orth only*: anchor links that were included only as orthologs; *Both orth and seq*: anchor links that were both included as orthologs and sequence similarity relationships; *Seq only*: anchor links that were included only as sequence similarity; *Pairs sharing func*: cross-species protein pairs that share at least one function. Species abbreviations are identical to Table S1.

**Table S3.**
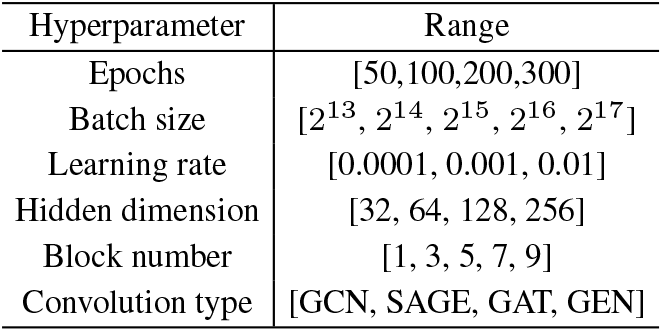
The search space of hyperparameters for training GraNA. GraNA is trained on train set and validated on valid set. The final combination of hyperparameters is determined based on GraNA’s performance on the valid set.

**Table S4.** AUROC of GraNA and baseline methods for predicting network alignment across species. For each dataset, we reported the AUROC values averaged over five independent train/test data splits. The abbreviations are identical to Table S1.

| Method | sce-spo | hsa-sce | hsa-mmu | hsa-cel | hsa-dme |
| --- | --- | --- | --- | --- | --- |
| MMseqs2 | 0.5057 | 0.5095 | 0.5102 | 0.5117 | 0.5101 |
| IsoRank | 0.5650 | 0.5179 | 0.5104 | 0.5143 | 0.5129 |
| MUNK-f | 0.5644 | 0.5819 | 0.5372 | 0.5111 | 0.5641 |
| MUNK-b | 0.5566 | 0.5772 | 0.5288 | 0.5079 | 0.5576 |
| TARA-TS | 0.6241 | 0.6384 | 0.6495 | 0.5848 | 0.6346 |
| TARA++ | 0.6270 | 0.6311 | 0.6533 | 0.5921 | 0.6372 |
| ETNA | 0.7045 | 0.6631 | 0.5805 | 0.5784 | 0.5891 |
| GraNA-o | 0.7707 | 0.6944 | 0.6568 | 0.6174 | 0.6367 |
| GraNA-s | 0.7681 | 0.6952 | 0.6473 | 0.6000 | 0.6287 |
| GraNA | **0.7865** | **0.7165** | **0.6755** | **0.6335** | **0.6506** |

**Table S5.** AUPRC of GraNA and baseline methods for predicting network alignment across species. For each dataset, we reported the AUPRC values averaged over five independent train/test data splits. The abbreviations are identical to Table S1.

| Method | sce-spo | hsa-sce | hsa-mmu | hsa-cel | hsa-dme |
| --- | --- | --- | --- | --- | --- |
| MMseqs2 | 0.0598 | 0.0547 | 0.0683 | 0.0635 | 0.0572 |
| IsoRank | 0.0770 | 0.0476 | 0.0628 | 0.0558 | 0.0512 |
| MUNK-f | 0.0748 | 0.0562 | 0.0675 | 0.0569 | 0.0578 |
| MUNK-b | 0.0740 | 0.0556 | 0.0661 | 0.0559 | 0.0564 |
| TARA-TS | 0.1019 | 0.0841 | 0.1140 | 0.0720 | 0.0842 |
| TARA++ | 0.0927 | 0.0756 | 0.1168 | 0.0783 | 0.0861 |
| ETNA | 0.1832 | 0.1053 | 0.0914 | 0.0720 | 0.0706 |
| GraNA-o | 0.2635 | 0.1258 | 0.1320 | 0.0931 | 0.0956 |
| GraNA-s | 0.2670 | 0.1336 | 0.1359 | 0.0970 | 0.1010 |
| GraNA | **0.2892** | **0.1511** | **0.1518** | **0.1078** | **0.1120** |

**Table S6.** Running time analysis of GraNA and baseline methods TARA, TARA-TS, and ETNA. The time needed for building topological features and training model were reported in minutes. Inference time could be neglected compared to feature-building and model-training time. The abbreviations are identical to Table S1.

| Method | Time | sce-spo | hsa-sce | hsa-mmu | hsa-cel | hsa-dme |
| --- | --- | --- | --- | --- | --- | --- |
| TARA | feature | 47 | 205 | 150 | 137 | 162 |
|  | train | <1 | 1 | 2 | <1 | 1 |
| TARA-TS | feature | <1 | 1 | 1 | 1 | 1 |
|  | train | <1 | 1 | 2 | <1 | 1 |
| ETNA | feature | <1 | 7 | 7 | 6 | 7 |
|  | train | <1 | <1 | <1 | <1 | <1 |
| GraNA | feature | <1 | 7 | 7 | 6 | 7 |
|  | train | 9 | 76 | 117 | 20 | 75 |

## 3 Supplementary Figures

**Fig. S1.**
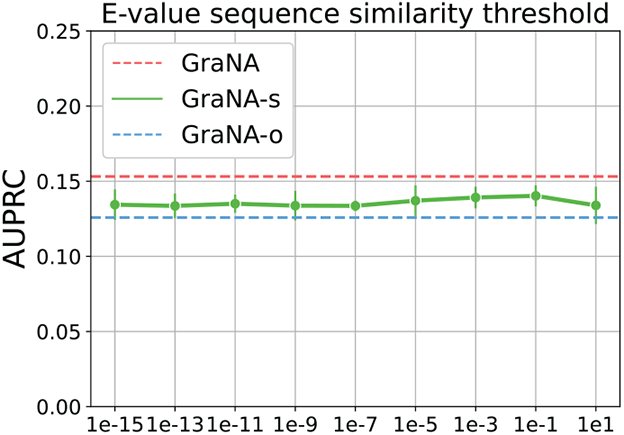
Impacts of E-value cutoff values used to identify sequence-similar protein pairs as anchor links. In GraNA, sequence similarity relationships are used as one type of anchor link. These sequence-similar protein pairs were identified by performing a sequence similarity search by MMseqs2 and selecting those pairs with an E-value smaller than a cutoff. We trained a GraNA variant that only used sequence similarity as anchor links (labeled as GraNA-s) and evaluated its AUPRC score of aligning the PPI networks of *H. sapiens* and *S. cerevisiae* when different E-value cutoffs were used. For reference, the AUPRC scores of GraNA-o and GraNA were shown. Since GraNA-o did not include sequence similarity as anchor links and GraNA used the default E-value cutoff of 10^−7^, their AUPRC scores were constant values in the figure.

**Fig. S2.**
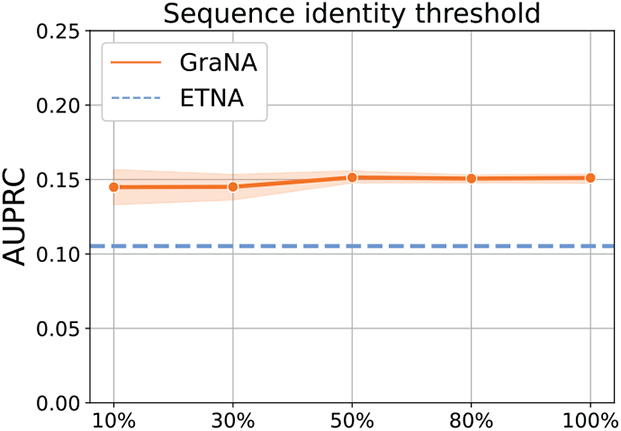
AUPRC of GraNA on data splits with different sequence identity thresholds. To validate GraNA’s effectiveness, we evaluate GraNA using harder data splits, which require that the train split and the test split are dissimilar in sequences. In practice, we fix the training sets and only filter test sets. Using MMseqs2 (Steinegger and Söding, 2017) to search the proteins in the test set that are under the sequence identity threshold, we constitute new test sets for each threshold. We select sequence identity thresholds 10%, 30%, 50%, 80%, and 100% (the original test split) and evaluate GraNA’s performance for each threshold on five independent data splits for *H. sapiens* and *S. cerevisiae*.

**Fig. S3.**
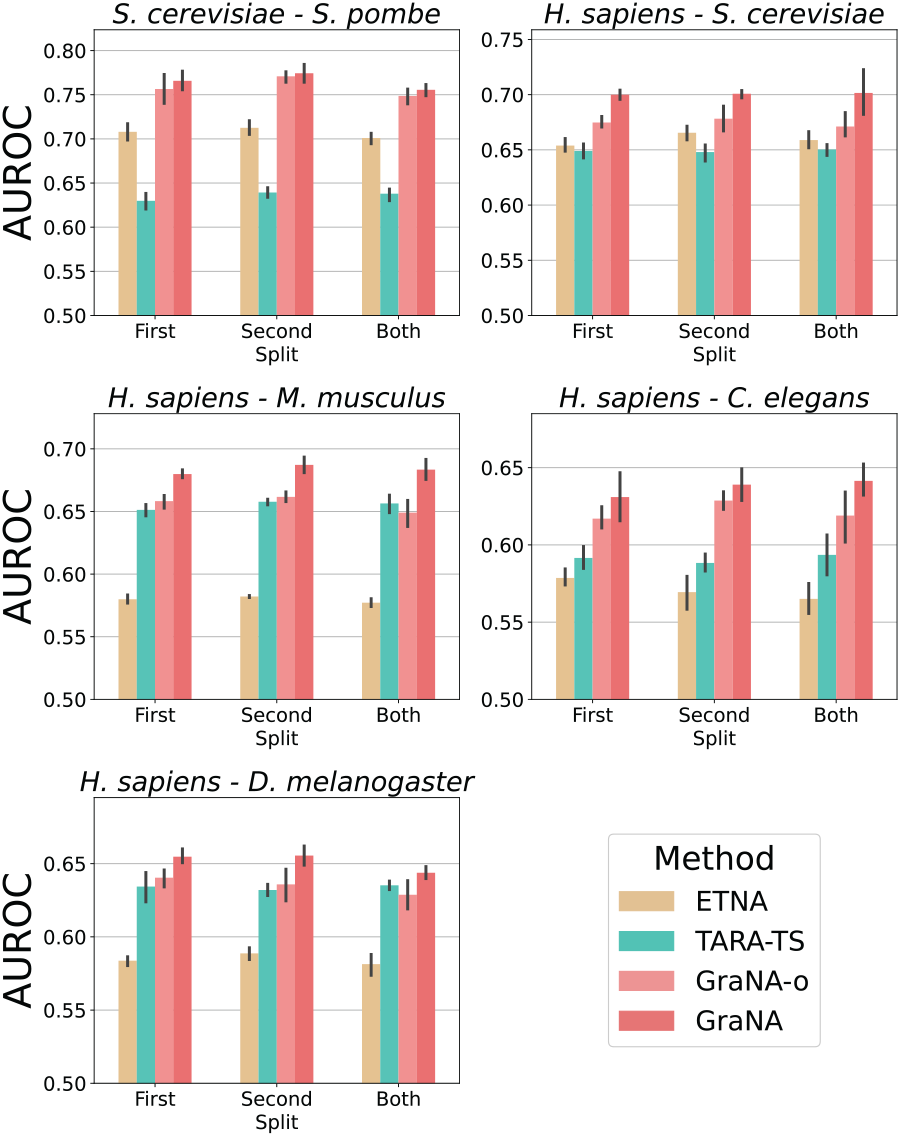
AUROC of network alignment prediction on sequence identity-based data splits. To further validate GraNA’s effectiveness under difficult data splits, we compared GraNA with the best unsupervised and supervised baselines (ETNA and TARA-TS) and a variant of GraNA (GraNA-o), that only uses orthologs as anchor links, on data splits that ensured proteins from the train split and the test split are dissimilar in terms of their sequence identity. Compared to the train/test splits in Fig. 2 where test proteins are ensured to not appear in the training set, here we create several more challenging train/test splits such that for the chosen species (the first species, the second species, or both species), its proteins in the test split must have sequence identity lower than 30% to its proteins in the train split. In our experiments, we iteratively sampled proteins and added those proteins together with their sequence-similar proteins (above 30% sequence identity) to the test set. The sequence identity is calculated by BLASTp (Camacho et al., 2009).

**Fig. S4.**
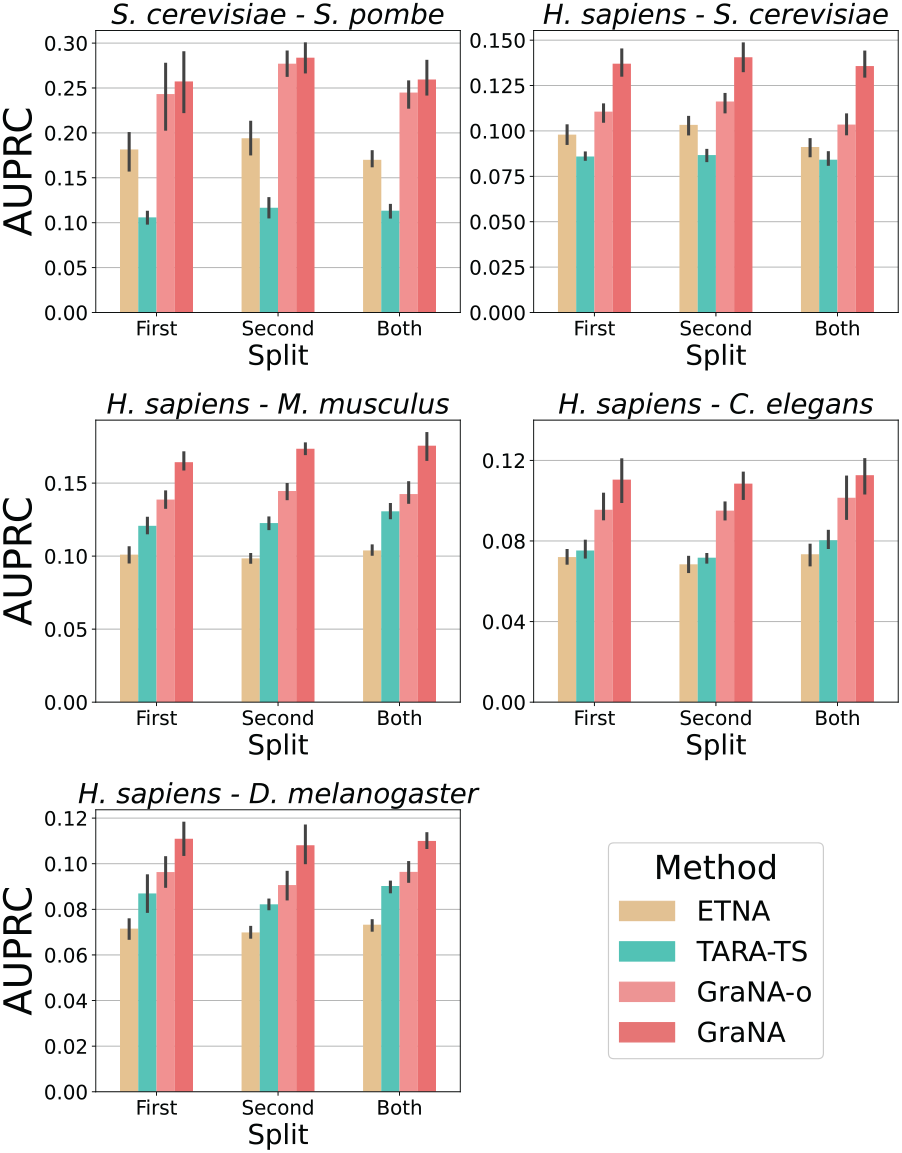
AUPRC of network alignment prediction on sequence identity-based data splits. To further validate GraNA’s effectiveness under difficult data splits, we compared GraNA with the best unsupervised and supervised baselines (ETNA and TARA-TS) and a variant of GraNA (GraNA-o), that only uses orthologs as anchor links, on data splits that ensured proteins from the train split and the test split are dissimilar in terms of their sequence identity. Compared to the train/test splits in Fig. 2 where test proteins are ensured to not appear in the training set, here we create several more challenging train/test splits such that for the chosen species (the first species, the second species, or both species), its proteins in the test split must have sequence identity lower than 30% to its proteins in the train split. In our experiments, we iteratively sampled proteins and added those proteins together with their sequence-similar proteins (above 30% sequence identity) to the test set. The sequence identity is calculated by BLASTp (Camacho et al., 2009).

**Fig. S5.**
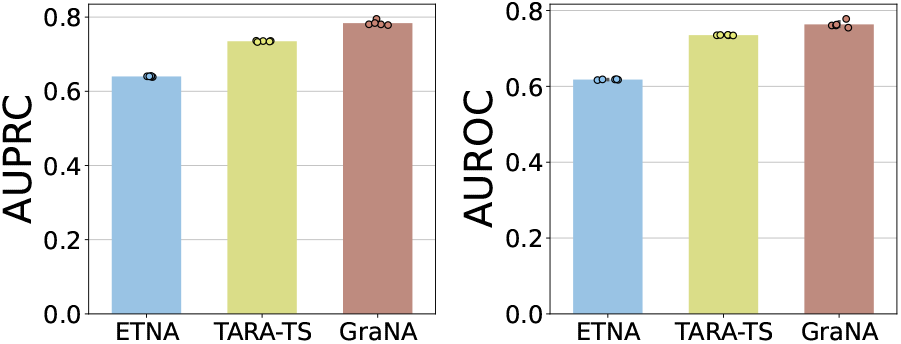
Network alignment performance on predicting newly discovered alignments between *H. sapiens*-*M. Musculus* based on known alignments. To further demonstrate GraNA’s potential for application, we compared GraNA with the best unsupervised and supervised baselines (ETNA and TARA-TS) on predicting the newly discovered alignments from GO (Consortium, 2004) (2022-12-04) that are not included in GO (Consortium, 2004) (2018-07-02). Following the method of generating the alignments in the benchmark dataset, we first create a slim set of GO terms from GO (2018-07-02) and then use it to generate new alignments in GO (2022-12-04), which contains 48% more functionally similar pairs. Supervised methods are trained on the supervision from 2018. All methods are evaluated on the dataset that includes all newly discovered alignments as positive samples and negative samples downsampled to an equal amount of positive samples. Experiments were repeated using five random seeds for negative sampling.

**Fig. S6.**
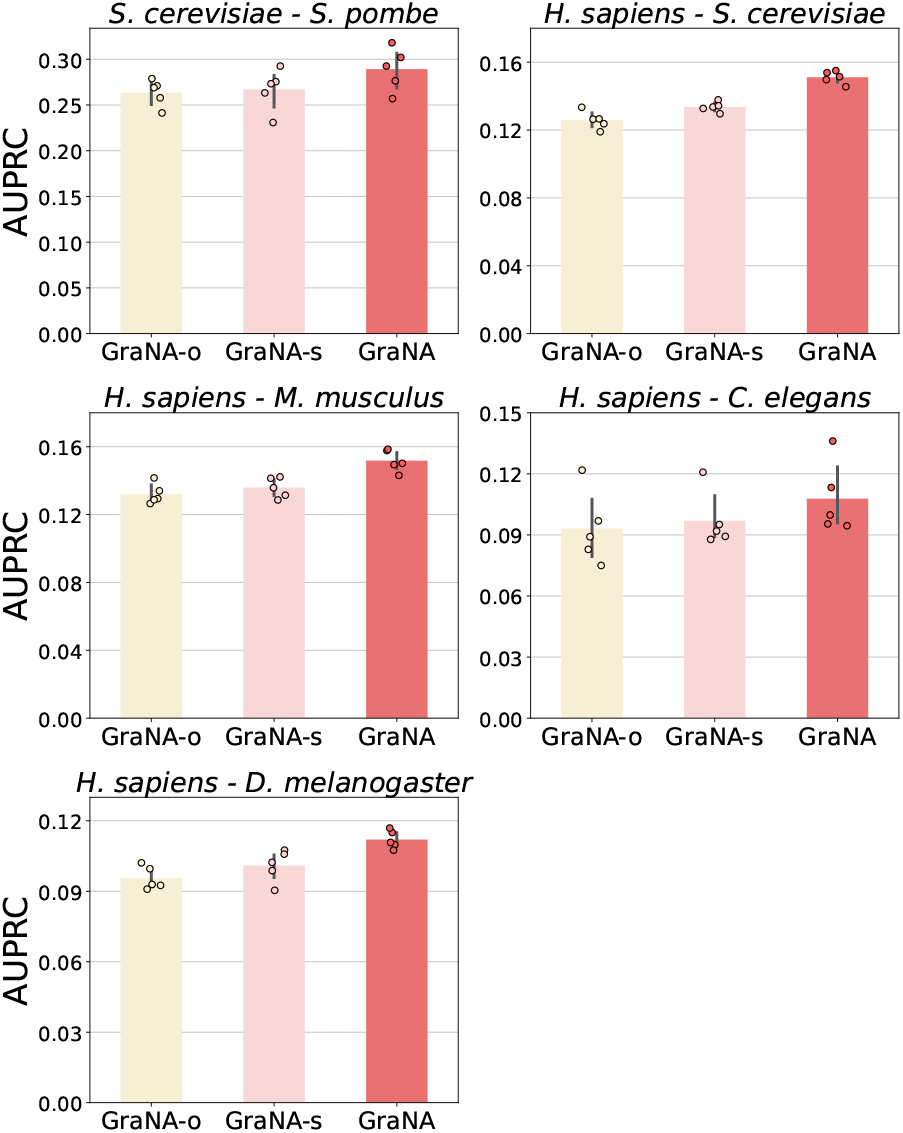
Network alignment performance of GraNA using different anchor links. We evaluated the performances of GraNA using only orthologs, only sequence similarity, and both orthologs and sequence similarity as anchor links for network alignment. Five pairs of PPIs (*S. cerevisiae*-*S. pombe, H. sapiens*-*S. cerevisiae, H. sapiens*-*M. Musculus, H. sapiens*-*C. elegans, H. sapiens*-*D. melanogaster*) are used for evaluation. AUPRC of five independent train/test data splits were reported.

**Fig. S7.**
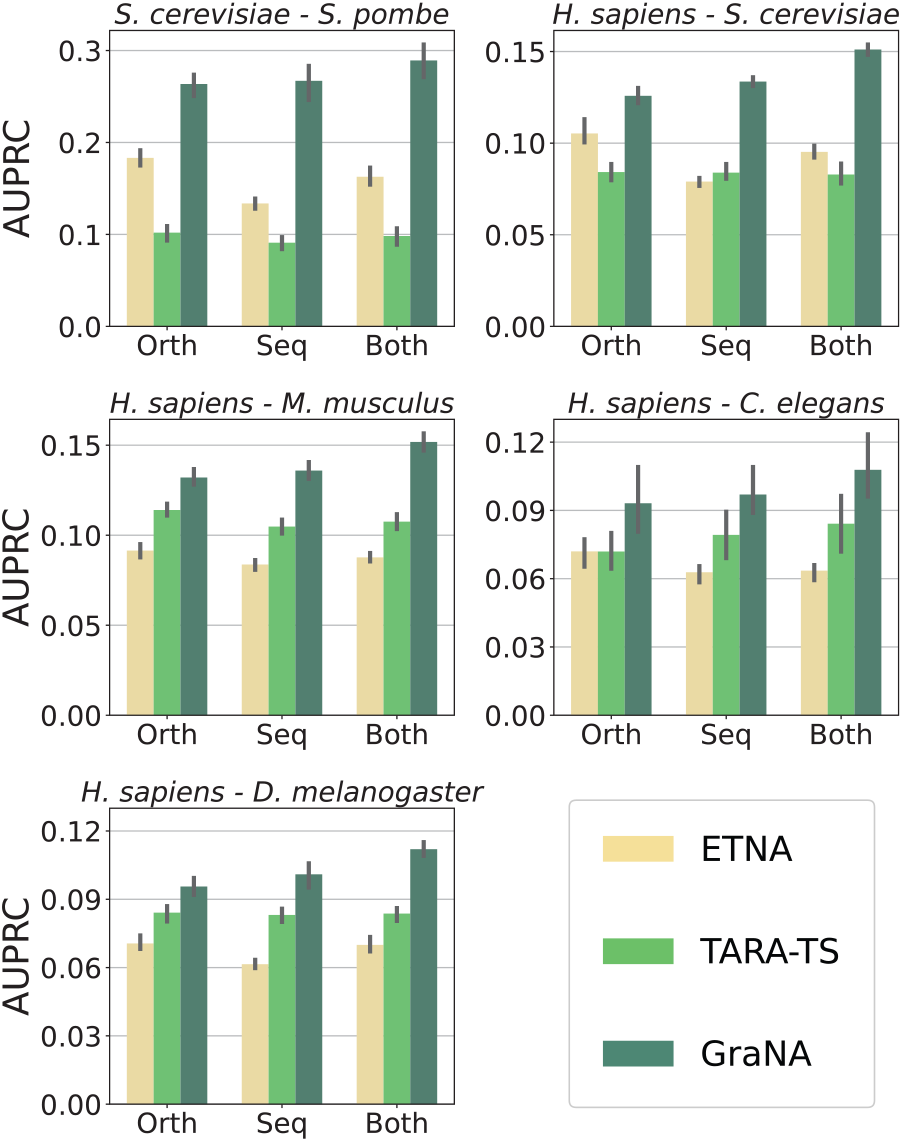
Evalutation of the ability to integrate heterogeneous anchor links. To validate GraNA’s ability to leverage orthology information and sequence similarity information at the same time, we compared GraNA with two of the best baselines, TARA-TS and ETNA, for network alignment using different anchor links. We used either orthologs, sequence similarity, or both orthologs and sequence similarity as anchor links for aligning five pairs of PPIs (*S. cerevisiae*-*S. pombe, H. sapiens*-*S. cerevisiae, H. sapiens*-*M. Musculus, H. sapiens*-*C. elegans, H. sapiens*-*D. melanogaster*), on five independent data splits. Abbreviations: orth: orthologs; seq: sequence similarity.

**Fig. S8.**
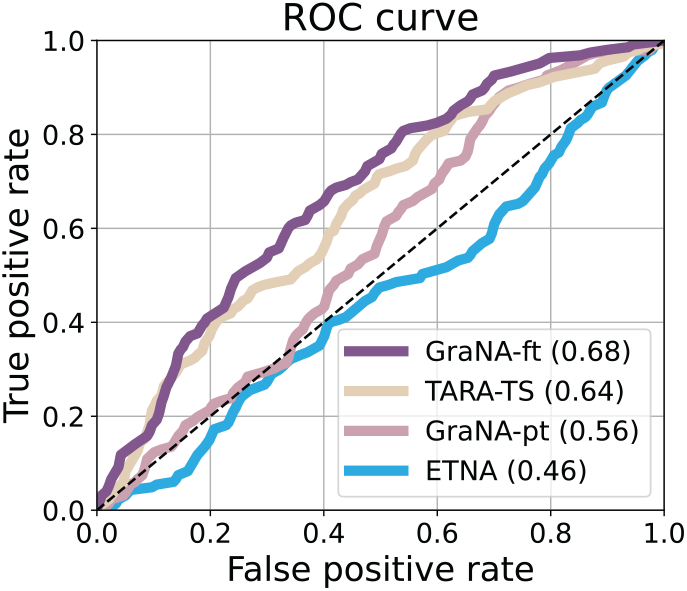
ROC curve of GraNA and baselines in the case study. We further included TARA-TS for comparison in predicting the replaceability of human genes with their yeast orthologs. TARA-TS is trained and evaluated on the dataset of experimental results by Kachroo et al. (Kachroo et al., 2015) via five-fold cross-validation.

**Fig. S9.**
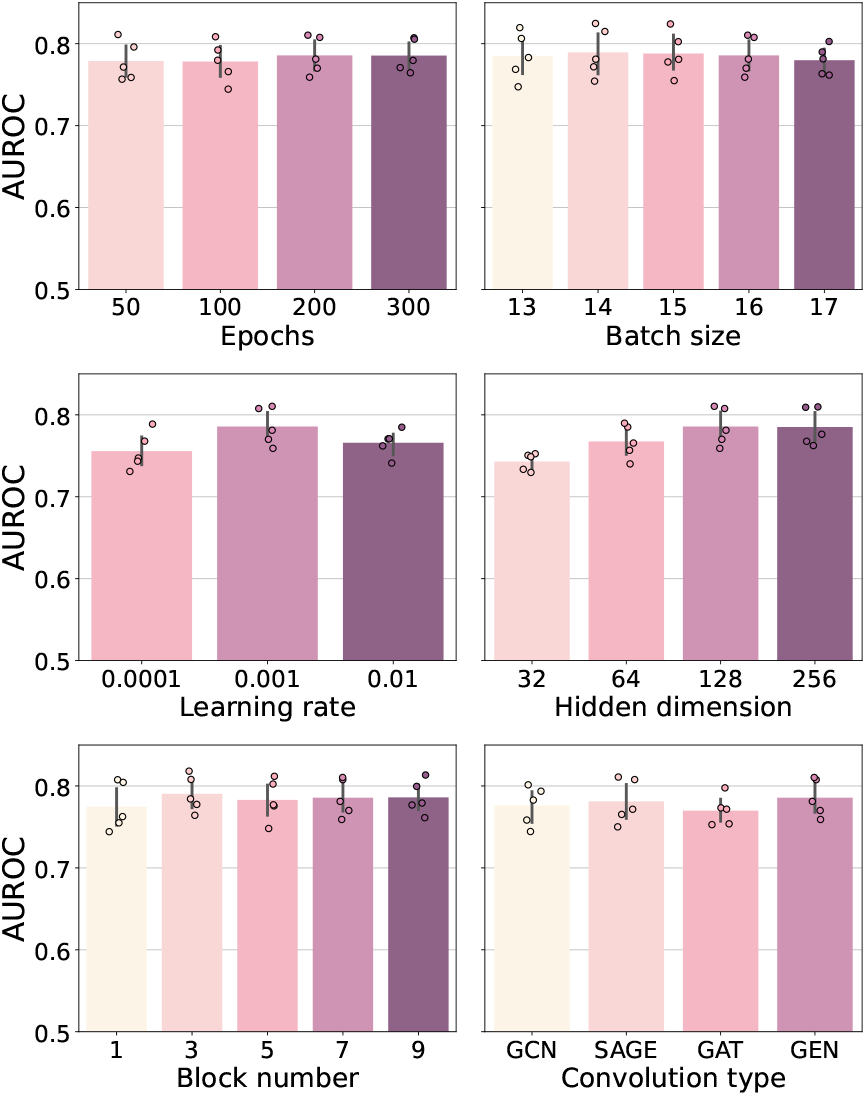
AUROC of GraNA for predicting the alignment between *S. cerevisiae* and *S. pombe* averaged over five independent data splits on valid set. While we are evaluating one type of hyperparameter, the other hyperparameters remain fixed. We evaluated in total 6 types of hyperparameters, including the total number of epochs, batch size, learning rate, hidden dimension, block number, and convolution type.

**Fig. S10.**
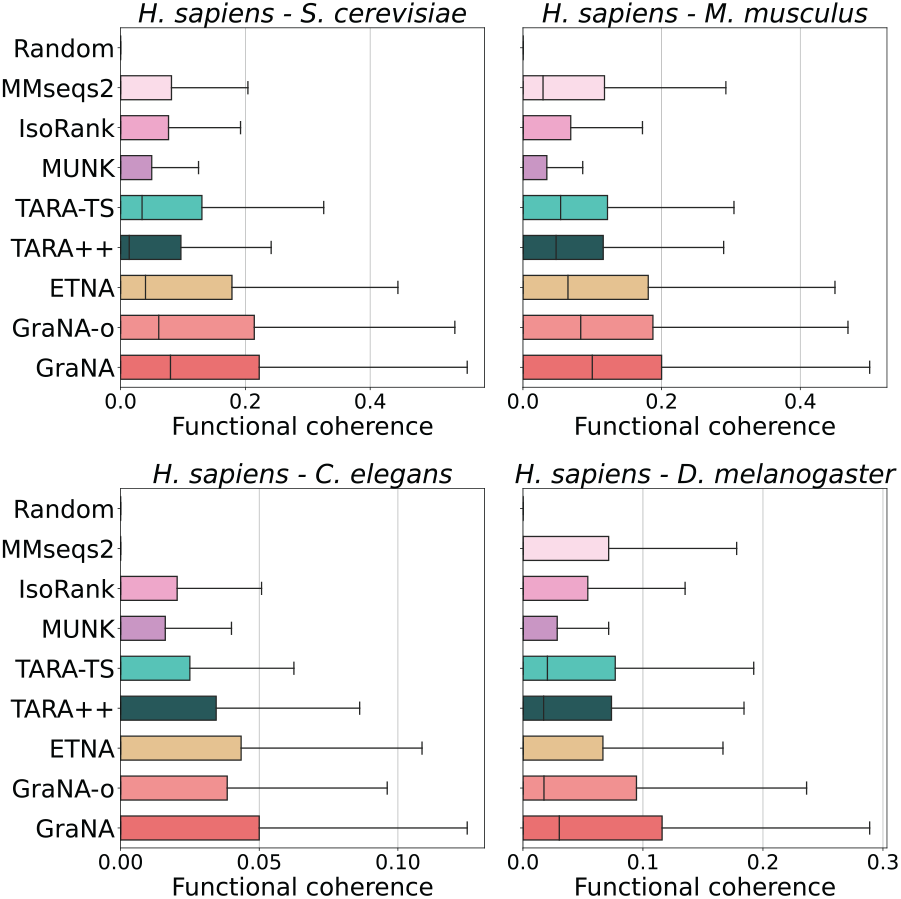
Functional coherence (FC) based on the network alignments produced by each method for four pairs of species (*H. sapiens*-*S. cerevisiae, H. sapiens*-*M. Musculus, H. sapiens*-*C. elegans, H. sapiens*-*D. melanogaster*). We chose the top 5,000 ranked protein pairs and transferred all the functional annotations of one protein in an aligned pair to predict the other protein’s function. The accuracy of the function prediction was evaluated by calculating the FC between the sets of the two aligned proteins. Unlike Jaccard index, FC only focuses on standardized GO terms (at a distance 5 to the root of the GO root) to avoid bias caused by terms from different levels of the GO hierarchy. Box plots showed the distribution of the FC of the top 5,000 aligned pairs for each method on five NA tasks under five random seeds.

**Fig. S11.**
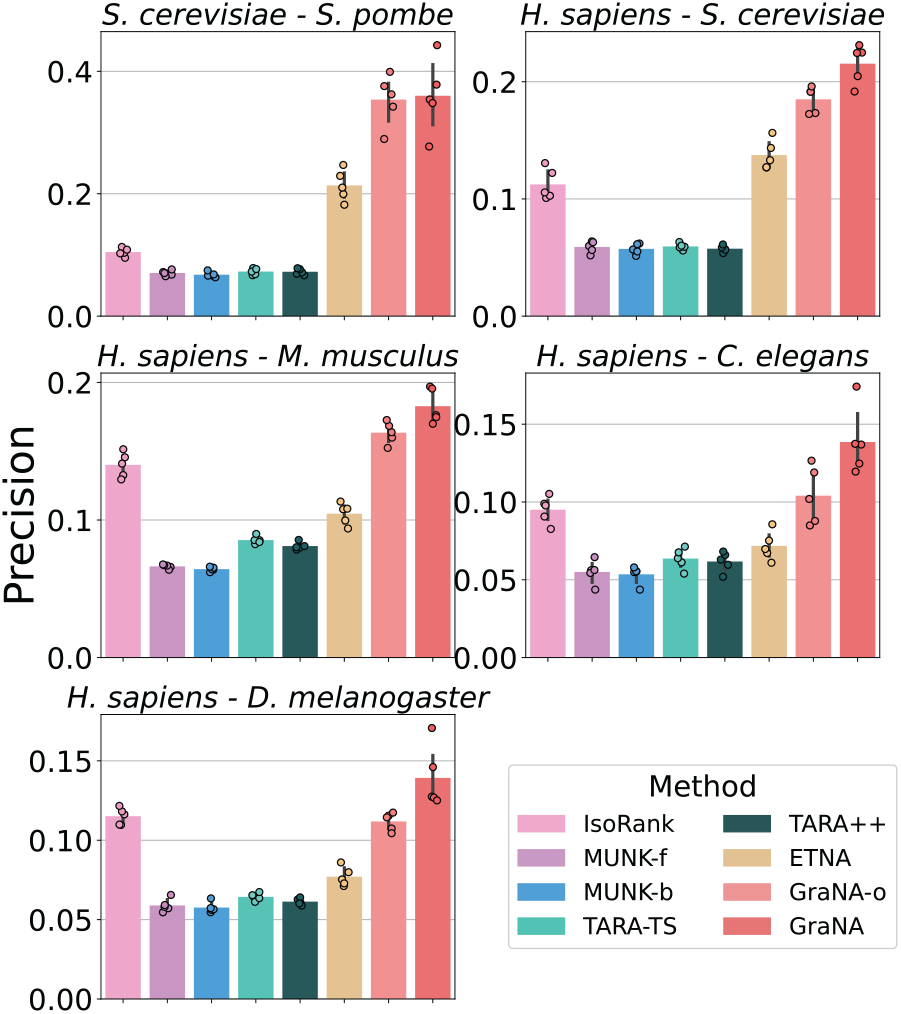
Precision of network alignment prediction. GraNA and other baselines were evaluated for aligning functionally similar proteins across five pairs of species, and we used precision as the metric. For GraNA’s predictions, we first selected the probability threshold maximizing the f1 score on the valid set and used this threshold to make final alignment predictions on the test set. GraNA-o is a variant of GraNA that only uses orthologs as anchor links whereas GraNA refers to the full model that uses both orthologs and sequence similarity as anchor links. As MUNK is not a bidirectional NA method, the performances of its forward and backward predictions were shown separately as MUNK-f and MUNK-b. Performances were evaluated using five independent train/test data splits.

**Fig. S12.**
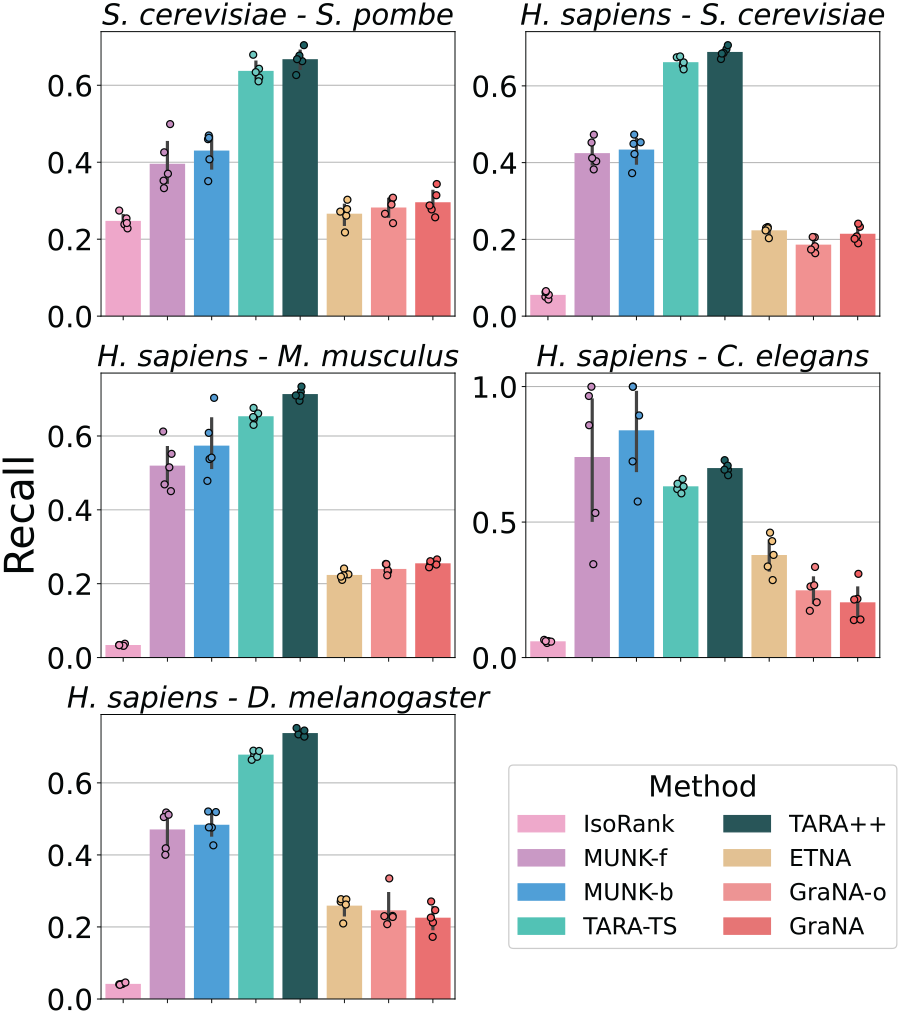
Recall of network alignment prediction. GraNA and other baselines were evaluated for aligning functionally similar proteins across five pairs of species, and we used recall as the metric. For GraNA’s predictions, we first selected the probability threshold maximizing the f1 score on the valid set and used this threshold to make final alignment predictions on the test set. GraNA-o is a variant of GraNA that only uses orthologs as anchor links whereas GraNA refers to the full model that uses both orthologs and sequence similarity as anchor links. As MUNK is not a bidirectional NA method, the performances of its forward and backward predictions were shown separately as MUNK-f and MUNK-b. Performances were evaluated using five independent train/test data splits.

**Fig. S13.**
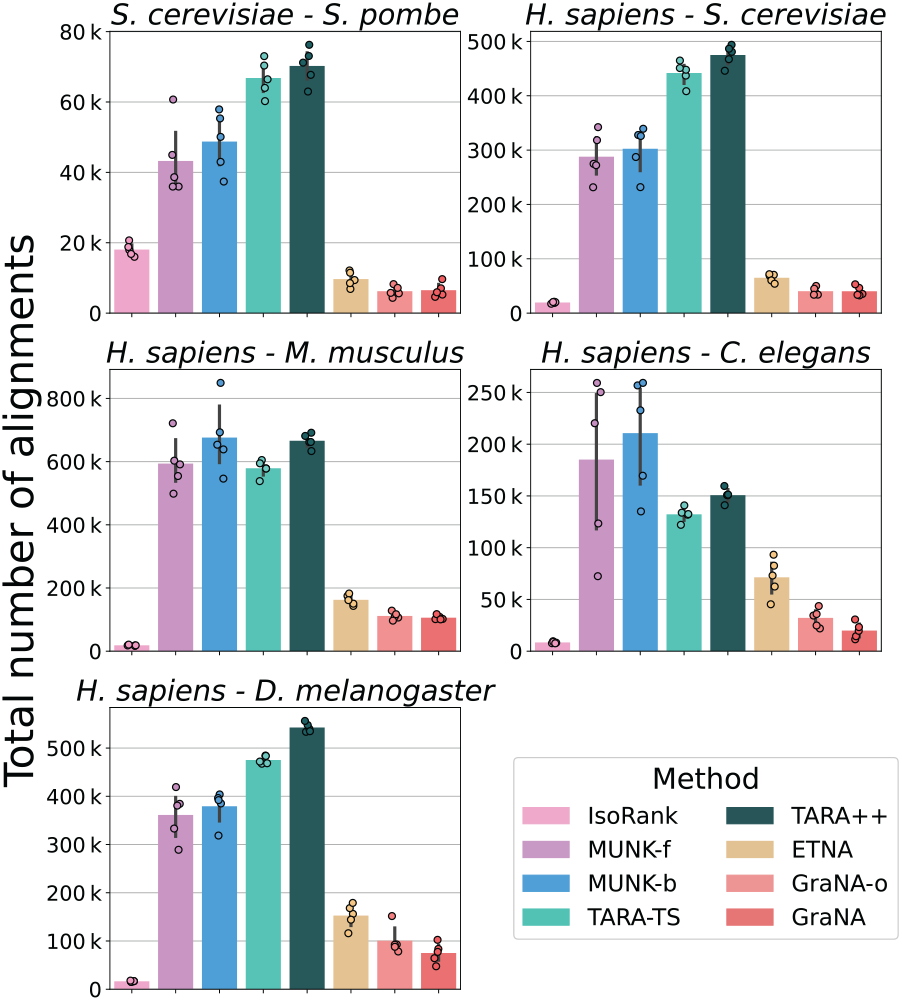
Total number of predicted network alignments. GraNA and other baselines were evaluated for aligning functionally similar proteins across five pairs of species, and we reported the total number of network alignments predicted by each method. For GraNA’s predictions, we first selected the probability threshold maximizing the f1 score on the valid set and used this threshold to make final alignment predictions on the test set. GraNA-o is a variant of GraNA that only uses orthologs as anchor links whereas GraNA refers to the full model that uses both orthologs and sequence similarity as anchor links. As MUNK is not a bidirectional NA method, the performances of its forward and backward predictions were shown separately as MUNK-f and MUNK-b. Performances were evaluated using five independent train/test data splits.

**Fig. S14.**
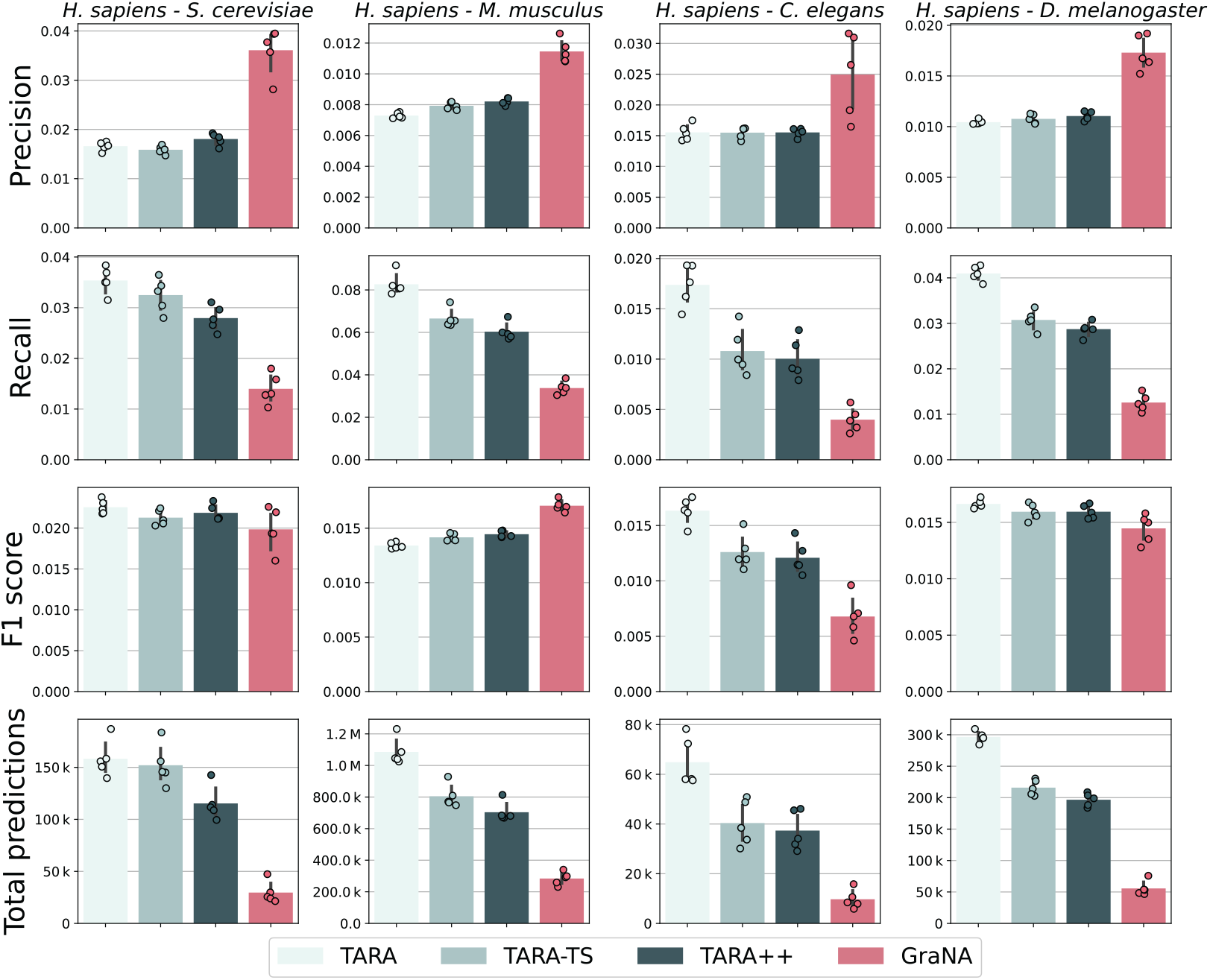
Performances of protein function prediction. We evaluated TARA, TARA-TS, TARA++, and GraNA in the context of cross-species protein function prediction in an established protein function prediction framework (Meng et al., 2016), and we used precision, recall, F1 score, and total number of predictions (protein-GO pairs) as metrics. The evaluation framework starts by performing network alignment prediction on a test set of protein pairs, and then it evaluates the functional predictions made based on the predicted network alignment via statistical tests. Consistent with other experiments in our manuscript, we only evaluated the methods on pairs of proteins that both have at least one alignment. As the data was unbalanced in the test set, we subsampled negative pairs of proteins to the number of positive pairs of proteins to construct a balanced test set. We restricted the function prediction only for GO terms from the slim set to avoid transferring general GO terms such as Biological Process. TARA++ prediction was the overlap of the predictions of TARA and TARA-TS. Performances were evaluated using five independent train/test data splits.

